# Human coronaviruses disassemble processing bodies

**DOI:** 10.1101/2020.11.08.372995

**Authors:** Mariel Kleer, Rory P. Mulloy, Carolyn-Ann Robinson, Danyel Evseev, Maxwell P. Bui-Marinos, Elizabeth L. Castle, Arinjay Banerjee, Samira Mubareka, Karen Mossman, Jennifer A. Corcoran

## Abstract

A dysregulated proinflammatory cytokine response is characteristic of severe coronavirus infections caused by SARS-CoV-2, yet our understanding of the underlying mechanism responsible for this imbalanced immune response remains incomplete. Processing bodies (PBs) are cytoplasmic membraneless ribonucleoprotein granules that control innate immune responses by mediating the constitutive decay or suppression of mRNA transcripts, including many that encode proinflammatory cytokines. PB formation promotes turnover or suppression of cytokine RNAs, whereas PB disassembly corresponds with the increased stability and/or translation of these cytokine RNAs. Many viruses cause PB disassembly, an event that can be viewed as a switch that rapidly relieves cytokine RNA repression and permits the infected cell to respond to viral infection. Prior to this report, no information was known about how human coronaviruses (hu CoVs) impacted PBs. Here, we show SARS-CoV-2 and the common cold hu CoVs, OC43 and 229E, induced PB loss. We screened a SARS-CoV-2 gene library and identified that expression of the viral nucleocapsid (N) protein from SARS-CoV-2 was sufficient to mediate PB disassembly. RNA fluorescent *in situ* hybridization revealed that N protein-mediated PB loss correlated with elevated RNA for PB-localized transcripts encoding TNF and IL-6. Ectopic expression of the N proteins from five other human coronaviruses (OC43, MERS, 229E, NL63 and SARS-CoV-1) did not cause significant PB disassembly, suggesting that this feature is unique to SARS-CoV-2 N protein. These data suggest that SARS-CoV-2-mediated PB disassembly contributes to enhanced proinflammatory cytokine production observed during severe SARS-CoV-2 infection.

## Introduction

Processing bodies (PBs) are membraneless ribonucleoprotein (RNP) granules found in the cytoplasm of all cells [1, 2] . PBs control cellular gene expression because they either degrade or sequester cellular RNA transcripts, preventing their translation into protein. PBs contain many enzymes required for mRNA turnover, including those needed for decapping (Dcp2 and co-factors Dcp1a and Edc4/Hedls) and decay of the RNA body (5’-3’ exonuclease Xrn1 and RNA helicase Rck/DDX6) and some components of the RNA-induced silencing complex [3, 4]. Not all coding RNAs are regulated by PBs, but those that are typically encode potent regulatory molecules like growth factors, pro-inflammatory cytokines, and angiogenic factors. One group of protein-coding mRNAs commonly found in PBs bear destabilizing AU-rich elements (AREs) in their 3’-untranslated regions (3’-UTRs) and include most proinflammatory cytokine transcripts [5–7]. These RNAs shuttle to PBs by virtue of interactions between the AU-rich element and RNA-binding proteins (RBPs) [3,8–11]. We and others showed that the presence of PBs correlates with increased turnover/suppression of ARE-mRNAs [7,12–15] . Conversely, when PBs are lost, constitutive ARE-mRNA suppression is reversed, and ARE-mRNA transcripts and/or their translation products accumulate. Therefore, PB disassembly can be viewed as a switch that permits cells to rapidly respond and translate ARE-containing proinflammatory cytokine RNA into molecules such as IL-6, IL-8, IL-1β, and TNF [5]. PBs provide an extra layer of post-transcriptional control enabling the cell to fine tune the production of potent molecules like proinflammatory cytokines.

Although PBs are constitutive, they are also dynamic, changing in size and number in response to different stimuli or stressors. This dynamic disassembly/assembly is possible because PBs behave as biomolecular condensates that form via liquid-liquid phase separation of proteins [16–19]. PBs form via sequential multivalent RNA-protein interactions, with a small group of proteins that contain regions of intrinsic disorder serving as the essential scaffold onto which additional proteins or RNA can be recruited as the PB matures [9,16,20–24]. Despite the recognition of PBs as dynamic entities, our understanding of the signals that induce PB disassembly remains incomplete. We and others have shown that stressors which activate the p38/MK2 MAP kinase pathway, as well as many virus infections elicit PB disassembly [12,13,15,25,26]. Disassembly can occur by a direct interaction between a viral protein(s) and a PB component that is subsequently re-localized to viral replication and transcription compartments (vRTCs) [27–29] or cleaved by viral proteases [30–32]. Viruses can also cause PB disassembly indirectly by activating p38/MK2 signaling [12,13,26] .

There are numerous reports of viral gene products that trigger PB disassembly, yet corresponding reports of viral gene products that stimulate PB formation are rare, suggesting that PBs possess direct antiviral function and their disassembly may favour viral replication in ways that we do not yet grasp [33, 34]. Even though other RNPs, such as stress granules, have emerged as important components of our antiviral defenses that contribute to sensing virus and triggering innate immune responses, evidence to support a direct antiviral role for PBs is less well established [34–37] . A direct-acting antiviral role has been defined for several PB-localized enzymes that impede viral replication (e.g. APOBEC3G, MOV10). However, in these cases, the mechanism of viral restriction was attributed to the enzymatic activity of the PB protein(s) whereas its localization to PBs was not deemed as significant [28,29,31,38–45]. Nonetheless, the disassembly of PBs by diverse viruses strongly suggests they negatively regulate virus replication. The reason viruses disassemble PBs may be to limit their antiviral activity; however, because PBs also control turnover/suppression of many proinflammatory cytokine transcripts, their disruption by viruses contributes to high proinflammatory cytokine levels, alerting immune cells to the infection.

The family *Coronaviridae* includes seven viruses that infect humans, including the four circulating ‘common cold’ coronaviruses (CoVs), CoV-OC43, CoV-229E, CoV-NL63, and CoV-HKU1 and three highly pathogenic viruses that cause severe disease in humans: MERS-CoV, SARS-CoV, and the recently emerged SARS-CoV-2 [46–51] . Severe COVID-19 is characterized by aberrant proinflammatory cytokine production, endothelial cell (EC) dysfunction and multiple organ involvement [52–64] . Even with intense study, we do not yet appreciate precisely how SARS-CoV-2 infection causes severe COVID-19 in some patients and mild disease in others though a mismanaged or delayed IFN response and an overactive cytokine response is thought to underlie severe outcomes [65–70] . Despite some contrasting reports, [71–73], what is clear is that SARS-CoV-2 proteins use a multitude of mechanisms to outcompete cellular antiviral responses [68,74–85].

To determine if SARS-CoV-2 and other CoVs interact with PBs to alter the cellular antiviral response, we performed an analysis of PBs after CoV infection. Prior to this research, no published literature was available on human CoVs and PBs, and only two previous reports mentioned PB dynamics after CoV infection. Murine hepatitis virus (MHV) was reported to increase PBs at early infection times, while transmissible gastroenteritis coronavirus (TGEV) infected cells displayed complete PB loss by 16 hours post infection [86, 87]. Observations that SARS-CoV-2 infection induced elevated levels of many PB-regulated cytokines, such as IL-6, IL-10, IL-1β and TNF [53,54,57,69,70] suggested that human CoVs like SARS-CoV-2 may reshape the cellular innate immune response in part by targeting PBs for disassembly. We now present the first evidence to show that three human CoVs, including SARS-CoV-2 trigger PB disassembly during infection. By screening a SARS-CoV-2 gene library, we identified that the nucleocapsid (N) protein was sufficient for PB disassembly. However, this feature is not common for all human coronavirus N proteins, as overexpression of MERS-CoV-N, OC43-N, 229E-N and NL63-N was insufficient to induce PB loss and expression of SARS-CoV-1 N protein displayed an intermediate phenotype. SARS-CoV-2 N protein expression was also sufficient to increase levels of inflammatory transcripts that localize to PBs and are known to be elevated during SARS-CoV-2 infection: IL-6 and TNF. Taken together, these results show that PBs are targeted for disassembly by human CoV infection, and that this phenotype may contribute in part to reshaping cytokine responses to SARS-CoV-2 infection.

## Results

### Infection with human coronaviruses causes PB loss

Endothelial cells (ECs) have emerged as playing a significant role in severe COVID; as sentinel immune cells they are important sources for many of the cytokines elevated in severe disease and are infected by SARS-CoV-2 *in vivo* [56,58–60,88]. However, others have shown that commercial primary human umbilical vein endothelial cells (HUVECs) require ectopic expression of the viral receptor, ACE2, to be susceptible to SARS-CoV-2 [89]. We recapitulated those findings and showed that after HUVECs were transduced with an ACE2-expressing lentivirus (HUVEC^ACE2^), they were permissive for SARS-CoV-2 (Wuhan-like ancestral Toronto isolate; TO-1) [90] (Fig S1A). To use HUVEC^ACE2^ for studies on PB dynamics, we confirmed that ACE2 ectopic expression had no effect on PB number in HUVECs (Fig S1B). Confirming this, we infected HUVEC^ACE2^ with SARS-CoV-2 (MOI=3) to determine if PBs were altered.

SARS-CoV-2 infected cells were identified by immunostaining for the viral nucleocapsid (N) while PBs were identified by immunostaining for two different PB resident proteins, the RNA helicase DDX6, and the decapping cofactor, Hedls. PBs, measured by staining for both markers, were absent in most SARS-CoV-2 infected HUVECs^ACE2^ by 24 hours post infection (Fig 1A-D). We quantified the loss of cytoplasmic puncta using a method described previously [91] and showed that by 24 hours post infection, SARS-CoV-2 infected cells displayed a significant reduction in PBs compared to mock-infected controls (Fig 1B, D).

**Figure 1.**
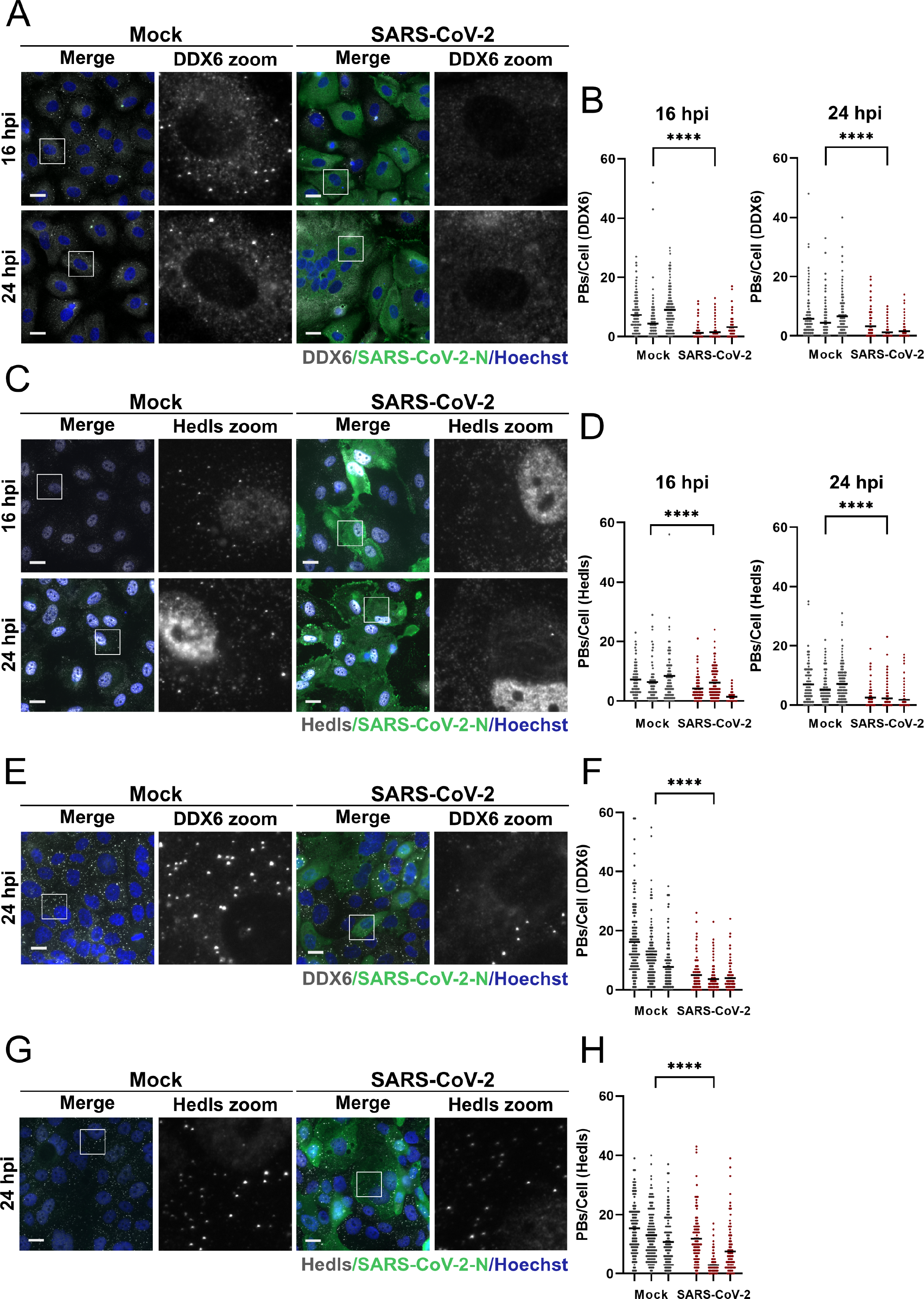
Processing bodies are absent in SARS-CoV-2 infected cells. **A-D**. HUVECs were transduced with recombinant lentiviruses expressing human ACE2 (HUVEC-ACE2), selected, and infected with SARS-CoV-2 TO-1 isolate at an MOI of 3 or a time-matched mock infection control. At 16 hpi or 24 hpi, cells were fixed and immunostained for SARS-CoV-2 N protein (green) and DDX6 (A, white) or Hedls (C, white). Nuclei were stained with Hoechst (blue). Representative images from one of three independent experiments are shown (A, C). PBs were quantified using CellProfiler by measuring DDX6 puncta (B) and Hedls puncta (D) in SARS-CoV-2-infected cells (thresholded by N protein staining) or in mock-infected cells. These data represent three independent biological replicates (*n=3*) with >80 cells measured per condition (mock and infected) per replicate. Each mock and infected replicate pair plotted independently (B, D); mean (****, *p* < 0.0001). E-H. Calu3 cells were infected with SARS-CoV-2 TO-1isolate (MOI =3) or time-matched mock-infected control. 24 hpi cells were fixed and immunostained for SARS-CoV-2 N (green), DDX6 (E; white) or Hedls (G; white). Nuclei were stained with Hoechst (blue). PBs were quantified using CellProfiler as in B and D (F, H). These data represent three independent biological replicates (*n=3*) with >130 cells measured per condition (mock and infected) per replicate. Representative images from one experiment of three are shown (E, G) and each mock and infected replicate pair plotted independently (F, H); mean (****, *p* < 0.0001). Statistics were performed using a Mann-Whitney rank-sum test. Scale bar = 20 µm.

To confirm that PBs were reduced by SARS-CoV-2 infection of naturally permissive cells derived from respiratory epithelium, we infected Calu-3 cells with SARS-CoV-2. Infected cells were identified by immunostaining for N protein 24 hours after infection and PBs were stained for DDX6 and Hedls. We observed PB loss in most but not all infected cells (Fig 1E-H). We quantified PB loss and showed that by 24 hours post infection, SARS-CoV-2 infected Calu-3 cells also displayed a significant reduction in PBs compared to mock-infected controls (Fig F, H). As the SARS-CoV-2 pandemic has progressed, new variants of concern (VOCs) have continued to emerge [92, 93]. To determine if VOCs also induced PB disassembly, we infected HUVEC^ACE2^ with SARS-CoV-2 VOCs Alpha, Beta, Gamma and Delta (MOI=2) and compared PB disassembly to an ancestral isolate of SARS-CoV-2 (Wuhan-like Toronto isolate; TO-1) (Fig 2). Infected cells were identified by immunostaining for N protein 24 hours after infection and PBs were identified using Hedls. Significant PB disassembly was observed for all VOCs, although we noted slightly less PB disassembly mediated by TO-1 compared to the experiments in Fig 1, likely due to the decrease in MOI between these two experiments (Fig 2).

**Figure 2.**
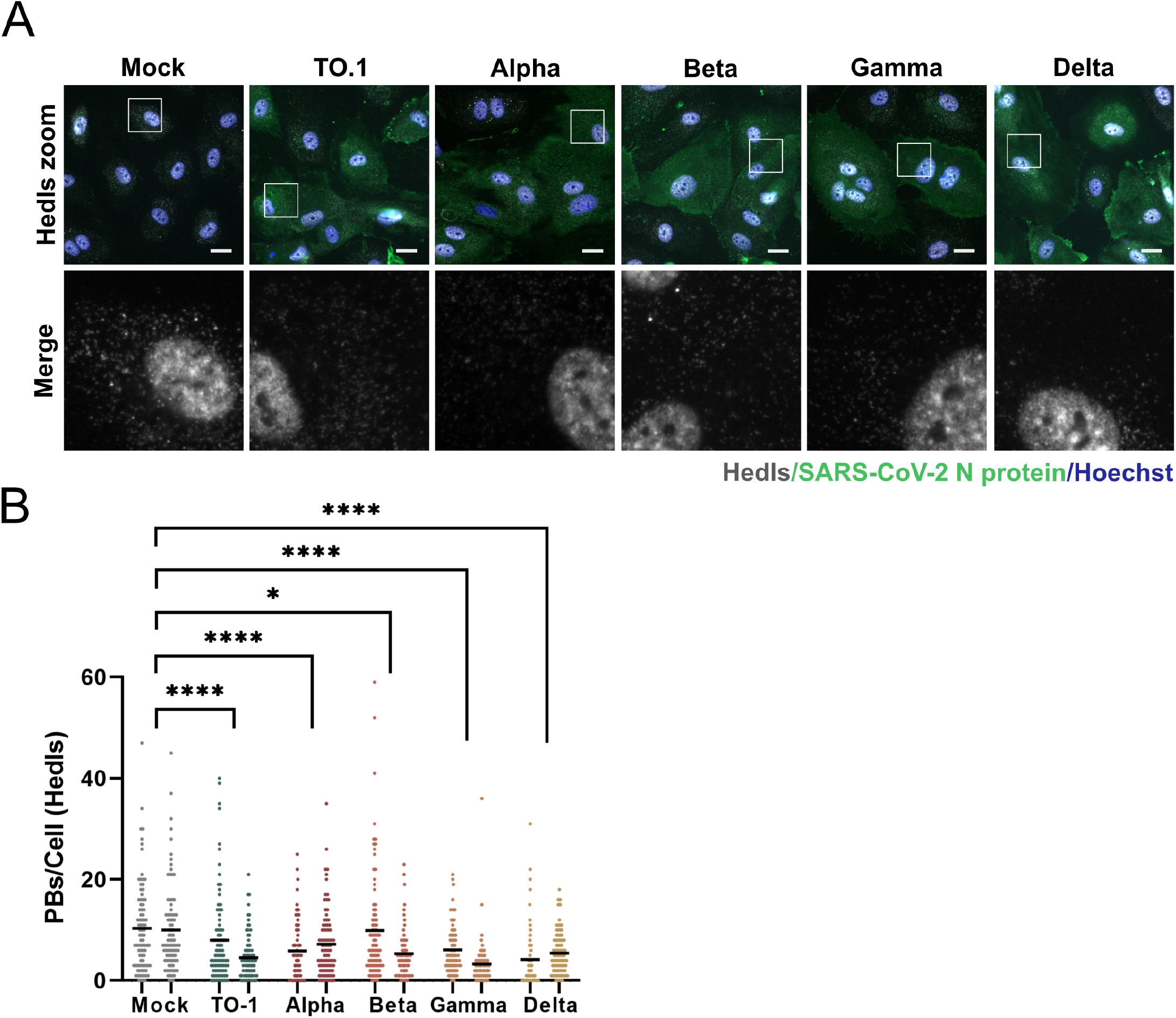
Processing bodies are absent in SARS-CoV-2 alpha, beta, gamma, and delta variant infected cells. **A.** HUVECs were transduced with recombinant lentiviruses expressing human ACE2 (HUVEC-ACE2), selected and infected with SARS-CoV-2 TO-1, alpha, beta, gamma, or delta isolates (MOI=2), or a mock infection control. 24 hpi cells were fixed and immunostained for SARS-CoV-2 N protein (green) and Hedls (white). Nuclei were stained with Hoechst (blue). Representative images from one of two independent experiments are shown. **B.** PBs were quantified as in Figure 1. These data represent two independent biological replicates (*n=2*) with >80 cells measured per condition (mock and infected) per replicate. Each mock and infected replicate pair plotted independently; mean. Statistics were performed using Kruskal-Wallis H test with Dunn’s correction (*, *p* < 0.0332; ****, *p* < 0.0001). Scale bar = 20 µM.

To determine if PBs were lost in response to infection with other human coronaviruses, we established infection models for the *Betacoronavirus*, OC43, and the *Alphacoronavirus*, 229E. We found HUVECs were permissive to both OC43 and 229E (Fig S2A-B). We then performed a time-course experiment wherein OC43-infected HUVECs were fixed at various times post infection and immunostained for the viral N protein and the PB-resident protein DDX6. We observed that PBs were largely absent in OC43 N protein-positive cells but present in mock-infected control cells (Fig 3A-B). 229E-infected HUVECs were stained for DDX6 to measure PBs and for dsRNA to denote infected cells due to a lack of commercially available antibodies for 229E. CoV infected cells are known to form an abundance of dsRNA due to viral replication and transcription from a positive-sense RNA genome making this a suitable marker for virally infected cells [94] . In parallel, we confirmed 229E infection by performing RT-qPCR for viral genomic and subgenomic RNA (Fig S2C-D). After 229E infection, we also found that PBs were significantly reduced (Fig 3C-D). Because of antibody incompatibility, we were unable to co-stain infected cells for the PB protein Hedls and OC43 N protein or dsRNA. In lieu of this, we performed additional OC43 and 229E infections and co-stained infected cells using antibodies for Hedls and DDX6 (Fig S3). We then performed additional quantification of PB loss using the Hedls marker to label PBs. These data also show robust PB loss in response to infection with OC43 and 229E (Fig S3).

**Figure 3.**
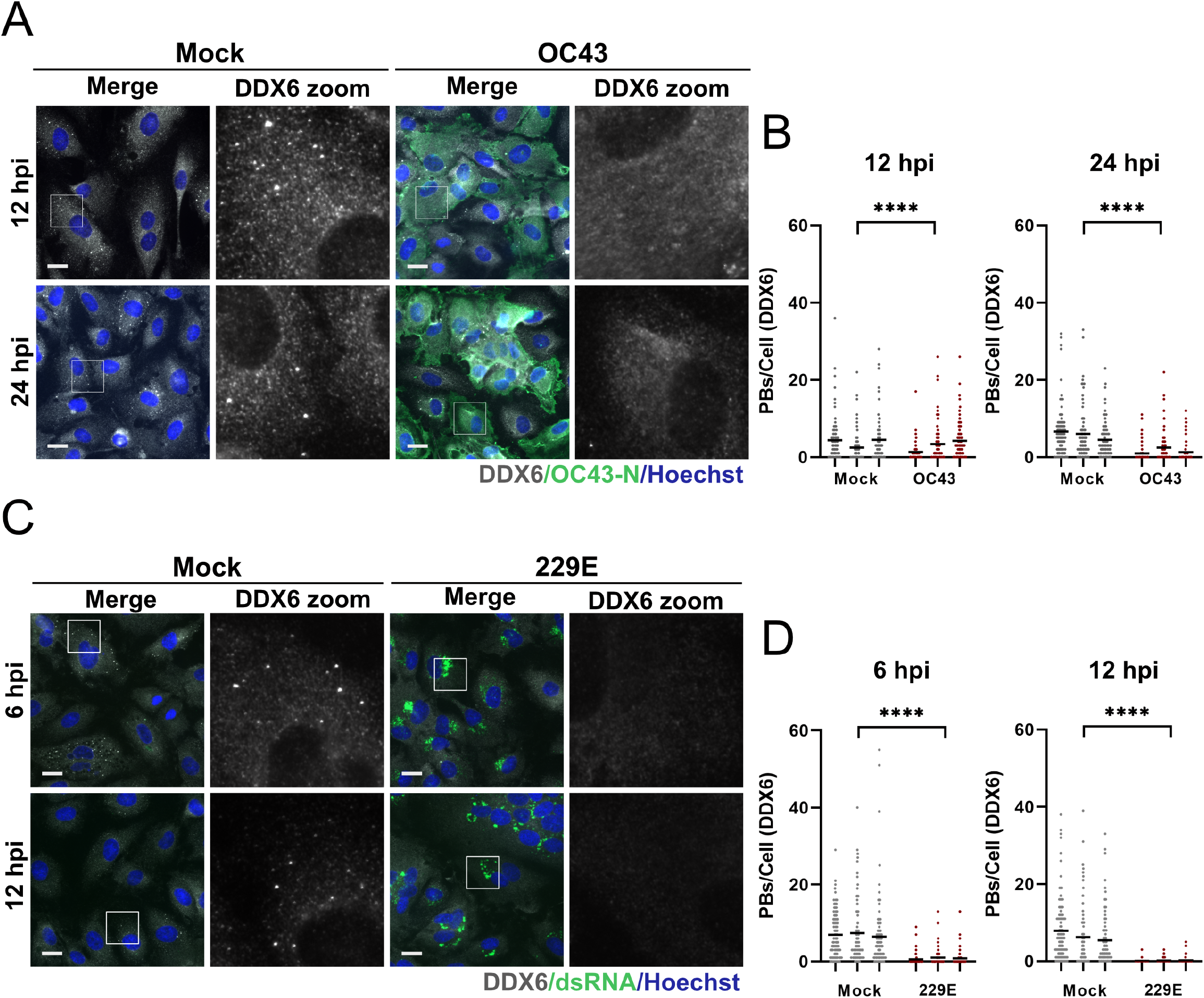
Processing bodies are absent in OC43 and 229E infected cells. **A-B.** HUVECs were infected with OC43 (TCID_50_ = 2 x 10^4^) or mock-infected. Cells were fixed at 12 hpi or 24 hpi and immunostained for DDX6 (PBs; white) and OC43 N protein (green). Nuclei were stained with Hoechst (blue). These data represent three independent biological replicates (*n=3*) with >80 cells measured per condition (mock and infected) per replicate. Representative images from one of three experiments are shown. DDX6 puncta in mock or OC43-infected cells were quantified as in Figure 1. Each mock and infected replicate pair plotted independently; mean. **C-D.** HUVECs were infected with 229E (TCID_50_ = 2.4 x 10^3^) or mock-infected. Cells were fixed at 6 hpi or 12 hpi and immunostained for DDX6 (PBs; white) and double-stranded RNA (proxy for 229E infection; green). Nuclei were stained with Hoechst (blue). These data represent three independent biological replicates (*n=3*) with >80 cells measured per condition (mock and infected) per replicate. Representative images from one of three experiments are shown. DDX6 puncta in mock or 229E-infected (dsRNA+) cells were quantified as in Figure 1. Each mock and infected replicate pair plotted independently; mean. Statistics were performed using a Mann-Whitney rank-sum test (****, *p* < 0.0001). Scale bar = 20 µM.

PBs will disassemble if key scaffolding proteins are lost; these include the RNA helicase DDX6, the translation suppressor 4E-T, the decapping cofactors Hedls/EDC4 and DCP1A, and the scaffolding molecule Lsm14A [95] . To elucidate if CoV-infected cells displayed decreased steady-state levels of PB resident proteins, we immunoblotted infected cell lysates for PB proteins XRN1, DCP1A, or DDX6, and Hedls (Fig 4) and quantified protein levels by densitometry (Fig S4). SARS-CoV-2 infection of HUVEC^ACE2^ cells did not alter steady-state levels of these proteins compared to uninfected cells (Fig 4A). OC43-infected HUVECs displayed comparable levels of XRN1, DCP1A, and Hedls relative to uninfected cells; however, OC43 infection decreased steady-state levels of DDX6 at both 12 hpi but not significantly at 24 hpi (Fig 4B). 229E-infected HUVECs showed no detectible change in PB protein expression after infection compared to controls (Fig 4C).

**Figure 4.**
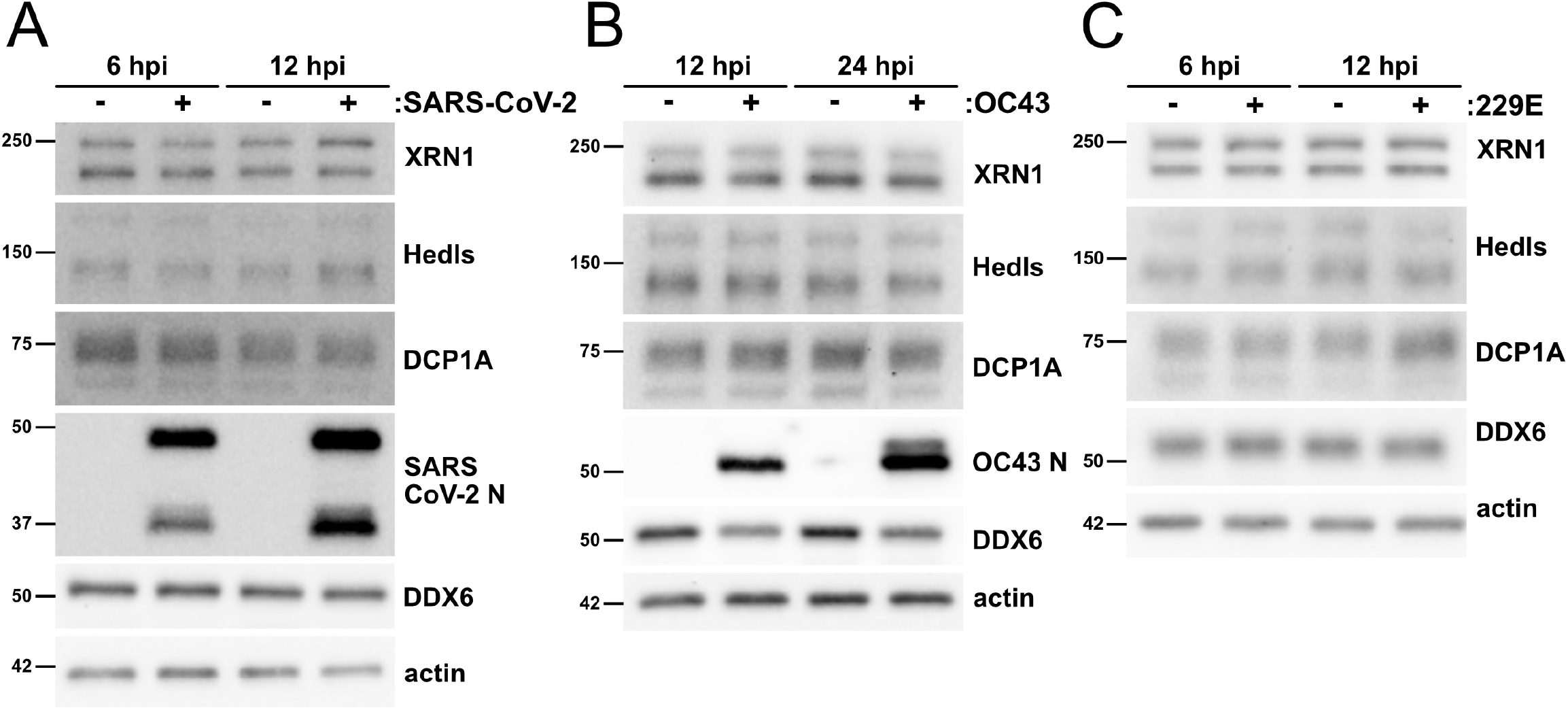
Coronavirus infection does not alter steady state levels of most processing body proteins. **A.** HUVECs were transduced with human ACE2, selected, and infected with SARS-CoV-2 TO-1 isolate (MOI=3). Cells were lysed at 6 and 12 hpi and immunoblotting was performed using XRN1, Hedls, DCP1A, DDX6, SARS-CoV-2 N, and β-actin specific antibodies. One representative experiment of two is shown. **B-C.** HUVECs were infected with OC43 (B, TCID_50_ = 2 x 10^4^) or 229E (C, TCID_50_ = 2.4 x 10^3^). Cells were lysed at 12 and 24 hpi (B, OC43) or 6 and 12 hpi (C, 229E). Immunoblotting was performed using XRN1, Hedls, DCP1A, DDX6, OC43 N protein (B only), and β-actin specific antibodies. One representative experiment of three is shown.

PBs are important sites for the post-transcriptional control of inflammatory cytokine transcripts containing AU-rich elements, and PB loss correlates with enhanced levels of some of these transcripts [7,12–15]. To determine if ARE-mRNAs are elevated, and therefore subject to regulation during CoV infection, we harvested total RNA from SARS-CoV-2-, OC43- or 229E-infected HUVECs and performed RT-qPCR for five ARE-containing cytokine transcripts, IL-6, CXCL8, COX-2, GM-CSF, and IL-1β (Fig 5A-C). Although we attempted to quantify TNF RNA as well, but the low copy number (high Ct value) of TNF RNA in mock control samples made quantification by RT-qPCR inaccurate and we were unable to measure the fold-increase of TNF RNA after CoV infection. We observed increased levels of three ARE-containing transcripts, IL-6, CXCL8, and COX-2, compared to uninfected cells, particularly in SARS-CoV-2 infected cells (Fig 5A). Taken together, these data indicate that infection with human coronaviruses including SARS-CoV-2 induces PB loss, and that some PB-regulated cytokine ARE-mRNAs are elevated during CoV infection.

**Figure 5.**
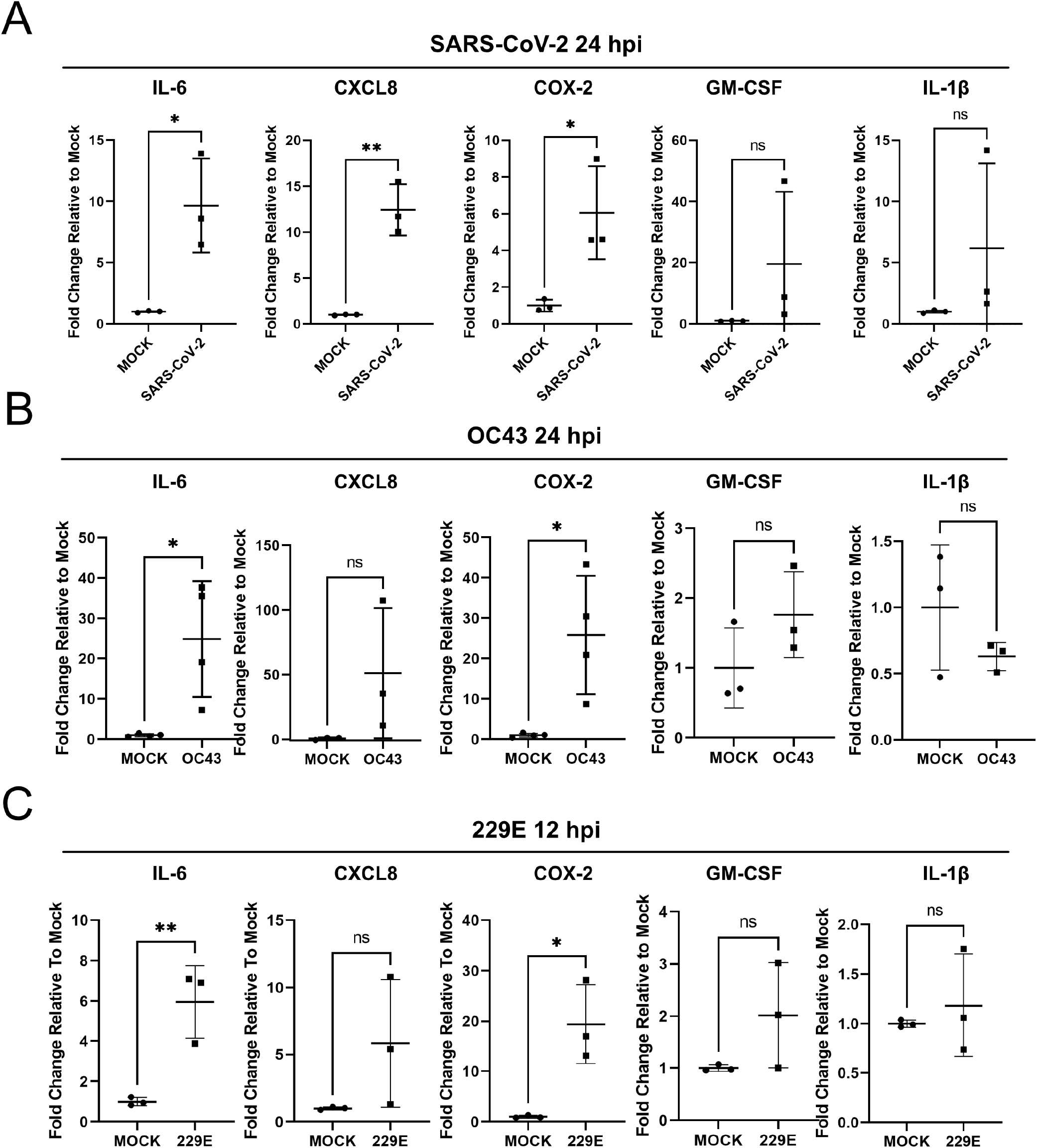
Steady state levels of selected ARE-mRNAs are elevated during coronavirus infection. **A.** HUVECs were transduced with recombinant lentiviruses expressing human ACE2, selected, and infected with SARS-CoV-2 TO isolate (MOI=3). RNA was harvested 24 hpi and RT-qPCR was performed using IL-6, CXCL8, COX-2, GM-CSF, IL-1β and HPRT (house-keeping gene) specific primers. Values are represented as fold change relative to mock-infection. *n*=3; mean ± SD (*, *p* < 0.05; **, *p*< 0.01; ns, nonsignificant). **B-C.** HUVECs were infected with OC43 (B, TCID_50_/mL = 3.5 x 10^4^) or 229E (C, TCID_50_/mL = 1 x 10^3.24^). RNA was harvested 24 and 12 hpi for OC43 (B) and 229E (C), respectively, and RT-qPCR was performed as in (A). Values are represented as fold change relative to mock-infection. *n*≥3; mean ± SD (*, *p* < 0.05; **, *p* < 0.01; ns, nonsignificant).

### A screen of SARS-CoV-2 genes reveals mediators of PB loss

The genome of SARS-CoV-2 is predicted to contain up to 14 open reading frames (ORFs). The two N-terminal ORFs (1a and 1ab) encode two large polyproteins which are processed by viral proteases into 16 non-structural proteins (nsp1-16) essential for viral genome replication and transcription [46] . The 3’ end of the SARS-CoV-2 genome is predicted to code for ORFs that are expressed from 9 subgenomic mRNAs [96]. Among these are the four structural proteins spike (S), envelope (E), membrane (M) and nucleocapsid (N) and up to 9 potential accessory proteins, not all of which have been validated as expressed in infected cells [96] . To test which SARS-CoV-2 gene product(s) was responsible for PB disassembly, we obtained a plasmid library of 27 SARS-CoV-2 genes from the Krogan lab; this library included 14 nsps (excluding nsp3 and nsp16), all structural (S, E, M, N) and candidate accessory genes (ORFs 3a, 3b, 6, 7a, 7b, 8, 9b, 9c, 10) [96]. We individually transfected each plasmid and immunostained for each of the SARS-CoV-2 proteins using antibodies to the Strep-tag II and for PBs using anti-DDX6 (Fig 6A, S5). Relative to control cells, many SARS-CoV-2 ORF transfected cells still displayed DDX6-positive PBs; however, expression of some SARS-CoV-2 genes reduced DDX6-positive puncta, including N and ORF7b (Fig 6A) to a similar or greater extent than our positive control, the KapB protein from Kaposi’s sarcoma-associated herpesvirus which we have previously shown causes PB disassembly (Fig 6B, C) [13,91,97,98]. We quantified the number of DDX6-positive PBs per cell for each transfection as in [91] and found that the average number of PBs per cell was reduced relative to our negative control after transfection of eight SARS-CoV-2 genes: nsp7, ORF7b, N, ORF9b, ORF3b, nsp6, nsp1, and nsp11 (Fig 6D-E). This quantification was performed in two different ways. In most cases, transfected cells were identified by co-staining for the Strep-tag II fused to each gene (Fig 6A, S5). In such cases, we quantified DDX6-positive puncta only in those cells that were transfected and not in bystander cells (Fig 6D, thresholded). These data identified three SARS-CoV-2 proteins that may cause PB loss in a cell autonomous manner: nsp7, ORF7b and N (Fig 6D). For the remaining transfections (nsp1, nsp5, nsp6, nsp11, nsp13, nsp14, ORF3b, ORF6, ORF9b, ORF9c) immunostaining for the Strep-tag II was not robust and we were unable to threshold our PB counts (Fig S5). In these samples, we quantified PBs in all cells of the monolayer (Fig 6E, unthresholded). These data identified five additional SARS-CoV-2 proteins from our screen that may cause PB loss: ORF9b, ORF3b, nsp1, nsp6 and nsp11 (Fig 6E). We verified the expression of all constructs, including low expressors (nsp1, nsp5, nsp6, nsp11, nsp13, nsp14, ORF3b, ORF6, ORF9b and ORF9c) by immunoblotting whole cell lysates harvested from parallel transfections (Fig 6F). We were unable to detect nsp4, ORF9c or ORF10 by immunoblotting; however, we did visualize expression of these proteins by immunostaining (Fig 6F, S5). We eliminated low confidence hits (nsp7, ORF9b) and low expressors (nsp6, nsp11) from further studies and proceeded with validation of the top four hits (ORF7b, N, ORF3b, nsp1).

**Figure 6.**
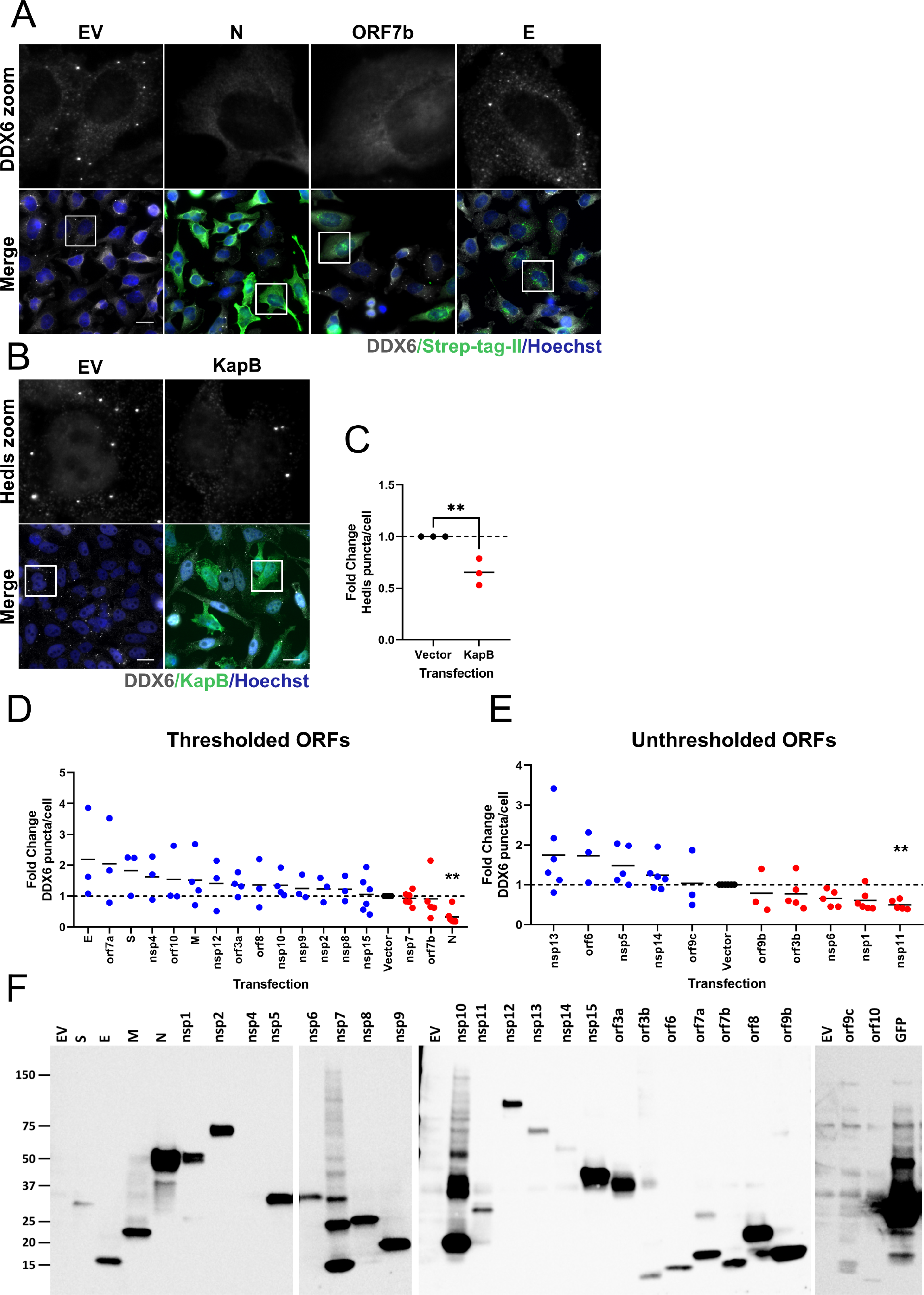
Identification of SARS-CoV-2 ORFs that mediate processing body loss. **A.** HeLa cells expressing GFP-Dcp1a were transfected with an empty vector (EV) or 2xStrep-tagged SARS-CoV-2 ORFs for 48 hours then fixed and immunostained for Strep-Tag-II (viral ORF; green) or DDX6 (PBs; white). Nuclei were stained with Hoechst (blue). Staining of select ORFs from one of three independent experiments are shown in A. **B.** As a positive control for PB disassembly, HeLa GFP-Dcp1a cells were transfected with EV or the Kaposi’s sarcoma-associated herpesvirus (KSHV) protein, KapB for 48 hours then fixed and immunostained for KapB (green) or Hedls (PBs; white). Nuclei were stained with Hoechst (blue); Scale bar = 20 μm. **C.** Hedls puncta were quantified as in Figure 1. These data represent three independent replicates (*n=3*). Statistics were performed using an unpaired T test; (**, *p* < 0.01). **D-E.** DDX6 puncta were quantified using CellProfiler. In D, SARS-CoV-2 ORF-expressing cells were thresholded by Strep-Tag-II staining intensity. The intensity threshold used was defined as two standard deviations above mean intensity in vector controls. Only DDX6 puncta in cells above this threshold were counted. In E, Strep-Tag-II staining could not be visualized above this threshold; therefore, DDX6 puncta in all cells were counted (unthresholded). Values are expressed as a fold-change difference normalized to the vector control (hashed line). These data represent three or more independent replicates (*n≥3*). A one-way ANOVA with a Dunnett’s post-hoc analysis was performed; bars represent SEM (**, *p* < 0.01). N=nucleocapsid protein, E=envelope protein, M=membrane protein, S=spike protein **F.** HeLa GFP-Dcp1a cells were transfected as in A. 48 hours post-transfection cells were lysed. Samples were resolved by SDS-PAGE on 4-15% gradient gels and immunoblotted with a Strep-Tag II antibody.

### The nucleocapsid protein of SARS-CoV-2 induces PB disassembly

We tested four top hits from our PB screen in more relevant endothelial cells as these cells can be infected, express inflammatory cytokines, and stain robustly for PBs. HUVECs were transduced with recombinant lentiviruses expressing N, nsp1, ORF3b, and ORF7b or empty vector control lentiviruses. We also included recombinant lentiviruses expressing nsp14 in this experiment because of its exoribonuclease activity and ability to diminish cellular translation and interferon responses [84]. Transduced cells were selected for transgene expression and then fixed and stained for the endogenous PB marker protein DDX6 and for the Strep-tag II on each of SARS-CoV-2 constructs. We observed robust staining of the viral nucleocapsid (N) protein in the transduced cell population (Fig 7A) but were unable to detect expression of nsp1, nsp14, ORF3b or ORF7b by immunostaining (Fig S6A). We quantified PB loss in the selected cells and observed decreased PB numbers in cell populations expressing N, nsp14, ORF3b and ORF7b; however, the most robust PB loss was induced in N-expressing cells, which displayed a significant reduction in PB numbers as well as strong immunostaining (Fig 7B, Fig S6). We were concerned that we could not detect the other four transgenes by immunostaining; therefore, we performed immunoblotting for the Strep-II tag on lysates from each transduced cell population (Fig S6C). Although we detected a strong band of ∼50 kDa at the predicted molecular weight for N, we were unable to detect bands for nsp14, ORF3b and ORF7b, while the most prominent band for nsp1 did not migrate at the predicted molecular weight of ∼20 kDa (Fig S6) [99, 100] . For these reasons, we decided to focus the remainder of our analysis on the N protein.

**Figure 7.**
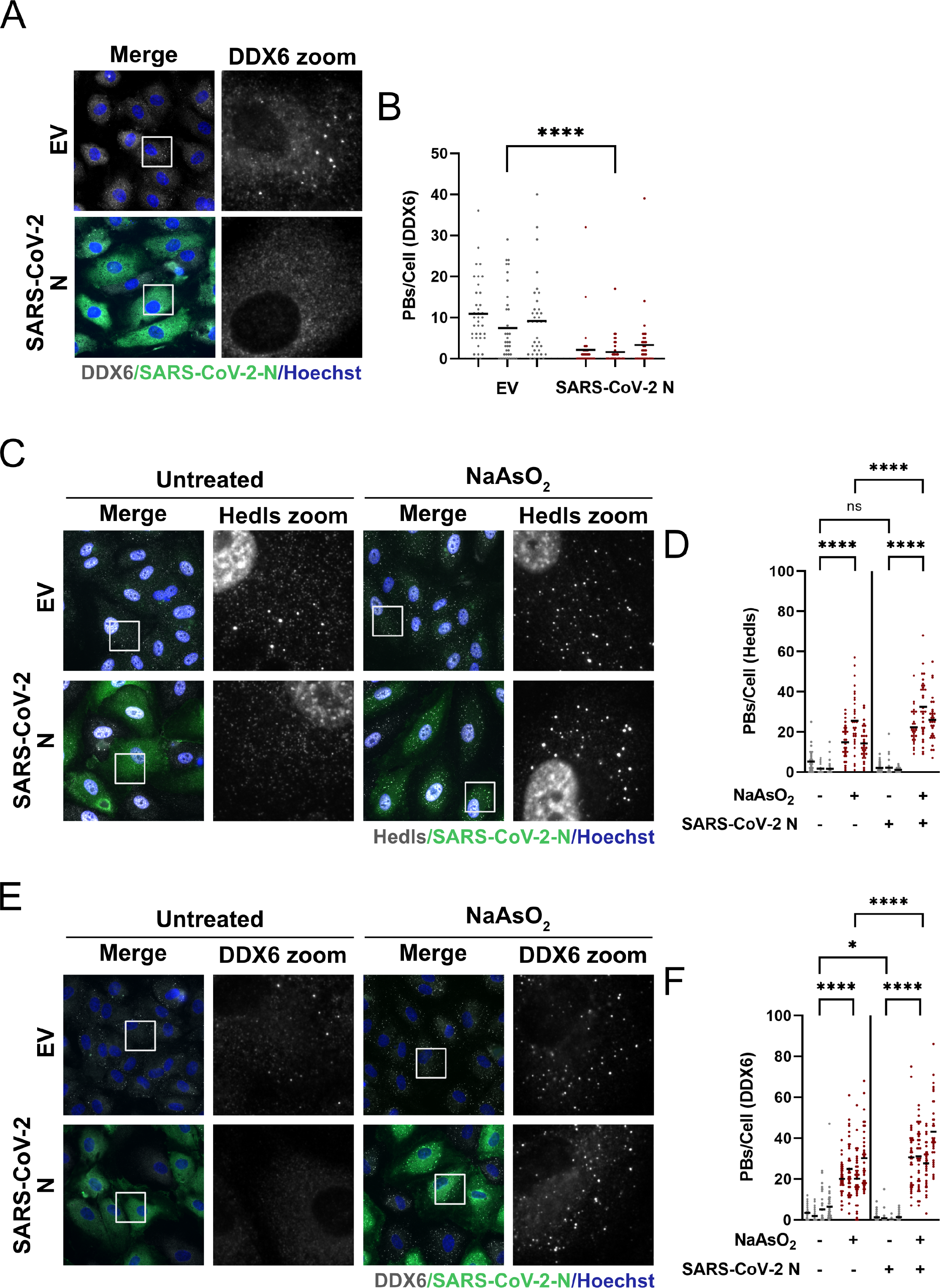
Ectopic expression of SARS-CoV-2 N elicits processing body disassembly. **A-B.** HUVECs were transduced with recombinant lentiviruses ectopically expressing 2xStrep-tagged SARS-CoV-2 N or empty vector control (EV) and selected with puromycin for 48 hours. Samples were fixed and immunostained for Strep-Tag-II (N; green) and DDX6 (PBs; white). Nuclei were stained with Hoechst (blue). Representative images from one of three independent experiments are shown (*n=3*). Scale bar = 20 μm. DDX6 puncta were quantified per field of view using CellProfiler as in Figure 1. >30 cells were measured per condition (EV and N) per replicate. Bars represent the mean. Mann-Whitney rank-sum test was performed (****, *p* < 0.0001). **C-F.** HUVECs were transduced and selected as in A, then treated with 0.25 mM sodium arsenite (NaAsO_2_) or a vehicle control (Untreated) for 30 min, fixed and immunostained for either Hedls (C, white) or DDX6 (D, white) and SARS-CoV-2 N (green). Nuclei were stained with Hoechst (blue). Scale bar = 20 μm. Representative images from one independent experiment of three are shown. These data represent three (C, D) or four (E, F) independent replicate s, with >30 cells measured per condition per replicate. Hedls puncta (D) and DDX6 puncta (F) were quantified as in Figure 1. Bars represent the mean. Statistics were performed using a two-way ANOVA with a Tukey’s multiple comparisons test (*, *p* < 0.0332; ****, *p* < 0.0001; ns, nonsignificant).

As PBs are dynamic RNP granules that undergo transient disassembly and assembly under basal conditions, we wanted to determine whether N-mediated PB loss was caused by enhanced disassembly of PBs or the prevention of PB *de novo* assembly. To determine this, we treated N-expressing HUVECs with sodium arsenite, a known inducer of PB assembly [101] and immunostained PBs for two different PB resident proteins, DDX6 and Hedls. Consistent with our previous observation, PB loss was observed after N expression in our untreated control (Fig 7C-F). However, in N-expressing cells treated with sodium arsenite, PB formation could still be observed, and was in fact increased, although the significance of this is unclear at present (Fig 7C-F). This means that N protein expression does not prevent PB formation, but instead triggers PB disassembly, which can be reversed after an appropriate stimulus. These data showed that N expression is sufficient to cause PB loss, and that the absence of PBs in N-expressing cells is a result of enhanced PB disassembly not a block in PB formation.

We showed that human CoVs OC43 and 229E also cause PB loss during infection (Fig 3); therefore, we were interested to determine if ectopic expression of nucleocapsid proteins from these or other human CoVs were sufficient to mediate PB disassembly. To test this, we transduced HUVECs with recombinant lentiviruses expressing the N protein from SARS-CoV-2 as well as N derived from Betacoronaviruses, MERS-CoV N-Flag and OC43 N. Expression of MERS-CoV and OC43 N proteins did not lead to significant PB loss compared to SARS-CoV-2-N (Fig 8A-B). We also tested two N proteins from human Alphacoronaviruses, 229E N-Flag and NL63 N-Flag. 229E N protein failed to induce significant PB loss compared to SARS-CoV-2 N, while unexpectedly, it appeared the expression of NL63 N protein increased PB numbers (Fig 8C-D). The significance of this increase is not yet clear. We also expressed the N protein from the more closely related Betacoronavirus, SARS-CoV-1, and observed SARS-CoV-1 N protein caused moderate PB loss, yet SARS-CoV-1 N protein-induced PB disassembly was not deemed significant by statistical analysis (Fig 8E-F). We collected protein lysates in parallel and performed immunoblotting to detect each CoV N protein (Fig 8G-I). We noted that CoV N proteins are not expressed equally, thus, we cannot fully discount expression level as the reason for the discrepancy in PB disassembly. This may be especially important for our analysis of SARS-CoV-1 N protein, which showed an intermediate but non-significant PB disassembly phenotype but was expressed at a lower level that SARS-CoV-2 N protein (Fig 8I).

**Figure 8.**
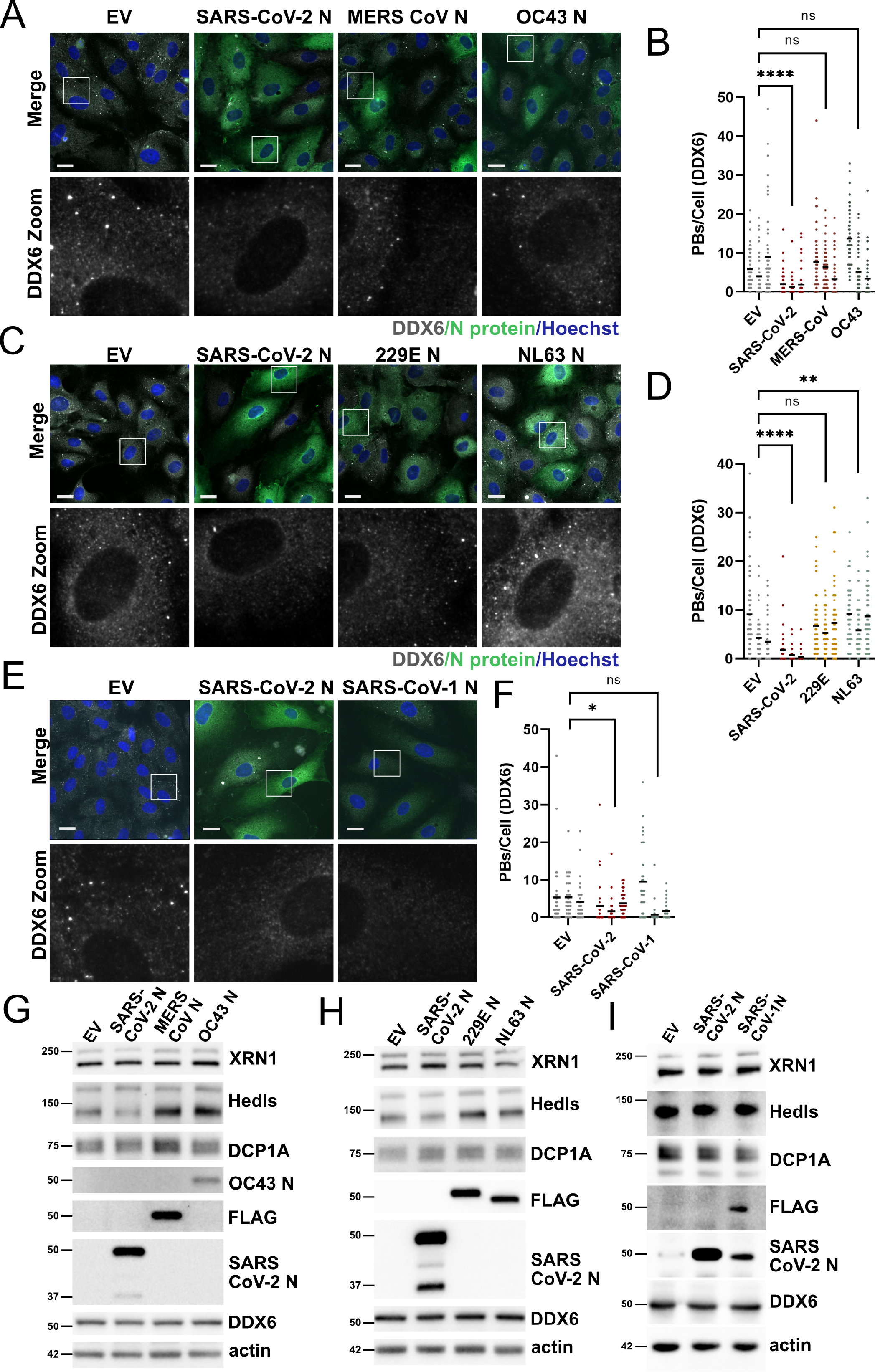
Processing body disassembly is not a common feature of all human coronavirus N proteins. **A-D.** HUVECs were transduced with recombinant lentiviruses ectopically expressing N protein from the betacoronaviruses MERS-CoV and OC43 (A-B) or N protein from alphacoronaviruses 229E and NL63 (C-D) or N protein from SARS-CoV-1 (E-F). A control lentiviral expressing an empty vector (EV) was used as a negative control and SARS-CoV-2 N protein expressing lentiviruses were used as a positive control in each experiment. Cells were selected, fixed and immunostained for DDX6 (PBs; white) and either authentic N protein or a FLAG tag (green). Nuclei were stained with Hoechst (blue). Scale bar = 20 µm. DDX6 puncta in EV or N-transduced cells were quantified using CellProfiler as in Figure 1. DDX6 puncta were quantified as in Figure 1. Representative images from one independent experiment of three are shown. These data represent three independent biological replicates (*n=3*) with >30 cells measured per condition (EV and N) per replicate. Each EV and N replicate pair plotted independently; mean. Statistics were performed using Kruskal-Wallis H test with Dunn’s correction (*, *p* < 0.0332; **, *p* < 0.0021; ****, *p* < 0.0001; ns, nonsignificant). **G-I.** HUVECs were transduced as above, protein lysate was harvested and immunoblotting was performed using XRN1, Hedls, DCP1A, DDX6, N protein or FLAG, and β-actin specific antibodies. One representative experiment of three is shown.

Immunoblotting of steady state levels of PB resident proteins after N protein overexpression showed that most PB proteins tested remained unchanged in the context of ectopic expression (Fig 8G-I, Fig S7A). The exception to this was the decapping factor, Hedls/EDC4, which was decreased after expression of SARS-CoV-2 N in one set of experiments but not in others, but increased after expression of N proteins from MERS and OC43 (Fig 8G-I, Fig S7A). The significance of this observation is also unclear.

To understand if PB disassembly correlates with changes to PB-regulated inflammatory cytokine transcripts in our system, we performed immunofluorescence-RNA fluorescent *in situ* hybridization (IF-FISH) to confirm the localization of inflammatory RNAs to PB foci. RNA FISH was performed for two PB-regulated cytokine transcripts that contain AREs, those encoding IL-6 and TNF, and for the GAPDH RNA, which does not contain AREs and is not expected to localize to PBs [3]. To achieve a better signal-to-noise ratio for detection, we used TNF to induce their transcription of IL-6 and TNF, which are lowly expressed in untreated cells.

First, we confirmed that TNF treatment alone did not significantly alter PB dynamics (Fig S8). We then stained PBs using our antibody to Hedls and co-stained with probes that bound GAPDH, TNF and IL-6 RNA transcripts. We repeatedly observed co-localization of both IL-6 and TNF RNA with PBs; in contrast, we observed extremely limited GAPDH co-localization with PBs (Fig 9A). IL-6 and TNF transcripts were also present at much lower levels than GAPDH RNA, consistent with the notion that ARE-containing transcripts are kept low by tight transcriptional control and constitutive decay in PBs [14]. We then stained N protein-transduced cells using our probes (Fig 9B). As expected, N protein expression induced loss of Hedls-positive PB foci (not shown). In N-expressing cells, the signal for IL-6 and TNF RNA redistributed from PB foci to the cytoplasm, and the probe intensity for both IL-6 and TNF was markedly increased compared to control cells which lacked N protein (Fig 9B). We quantified the change in FISH probe signal intensity per cell (Fig 9C). Compared to controls, N protein-expressing cells displayed strongly increased TNF probe signal intensity and increased IL-6 probe signal intensity, whereas GAPDH probe signal intensity was not increased (Fig 9C). These data support our hypothesis that PBs directly regulate proinflammatory cytokine transcripts and that these PB-regulated mRNAs increase upon PB disassembly.

**Figure 9.**
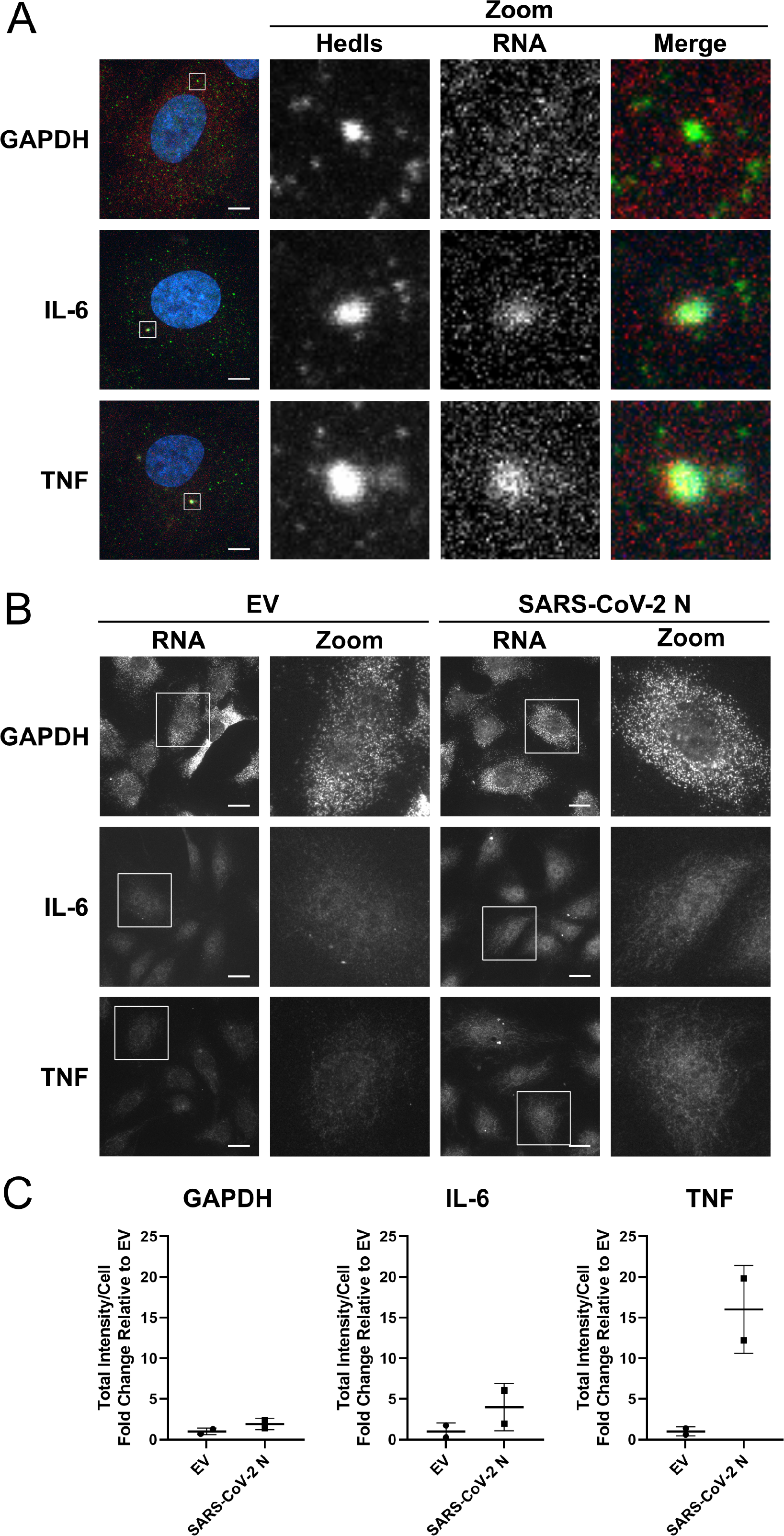
ARE-mRNAs colocalize with processing bodies. **A.** HUVECs were treated with 0.01 ng/L soluble TNF to increase transcription of ARE-containing cellular mRNAs for 24 hours. Cells were fixed and immunostained for Hedls prior to hybridization with Stellaris probes specific for GAPDH, IL-6 and TNF. Nuclei were stained with Dapi. Cells were imaged using a Zeiss LSM 880 confocal microscope with Airyscan using the 63x objective. At least three Z-stacks were acquired per condition (probe) per replicate and a maximal intensity projection (MIP) image is presented. One representative experiment of three is shown. Scale bar = 5 µm. **B-C.** HUVECs were transduced with recombinant lentiviruses ectopically expressing SARS-CoV-2 N or an EV control. Cells were selected, fixed and immunostained for Hedls prior to hybridization with Stellaris probes specific for GAPDH, IL-6 and TNF. Nuclei were stained with Dapi. Only probe-specific staining is displayed for simplicity. Scale bar = 20 μm. Representative images from one of two independent experiments are shown. Total signal intensity per cell was quantified using CellProfiler and is represented as fold-change relative to the intensity in the EV control.

We then asked if N protein from SARS-CoV-2 or other human CoVs would increase ARE-RNA transcript levels by RT-qPCR. We analyzed IL-6 and TNF transcript level, as well as two other ARE-containing RNAs, encoding CXCL8 and COX-2 which were elevated after infection with SARS-CoV-2 and OC43 (Fig 5). In uninduced ECs, the transcription of these mRNAs is minimal; for example, TNF mRNA could not be readily detected by RT-qPCR without transcriptional activation. Therefore, we treated control and N-expressing HUVECs with TNF to activate cytokine transcription and then assessed if N protein expression enhanced cytokine mRNA level post-transcriptionally. In the absence of TNF, no change of mRNA abundance was observed for any of the coronaviruses N proteins tested (Fig 10A-C, 0 hour no treatment).

**Figure 10.**
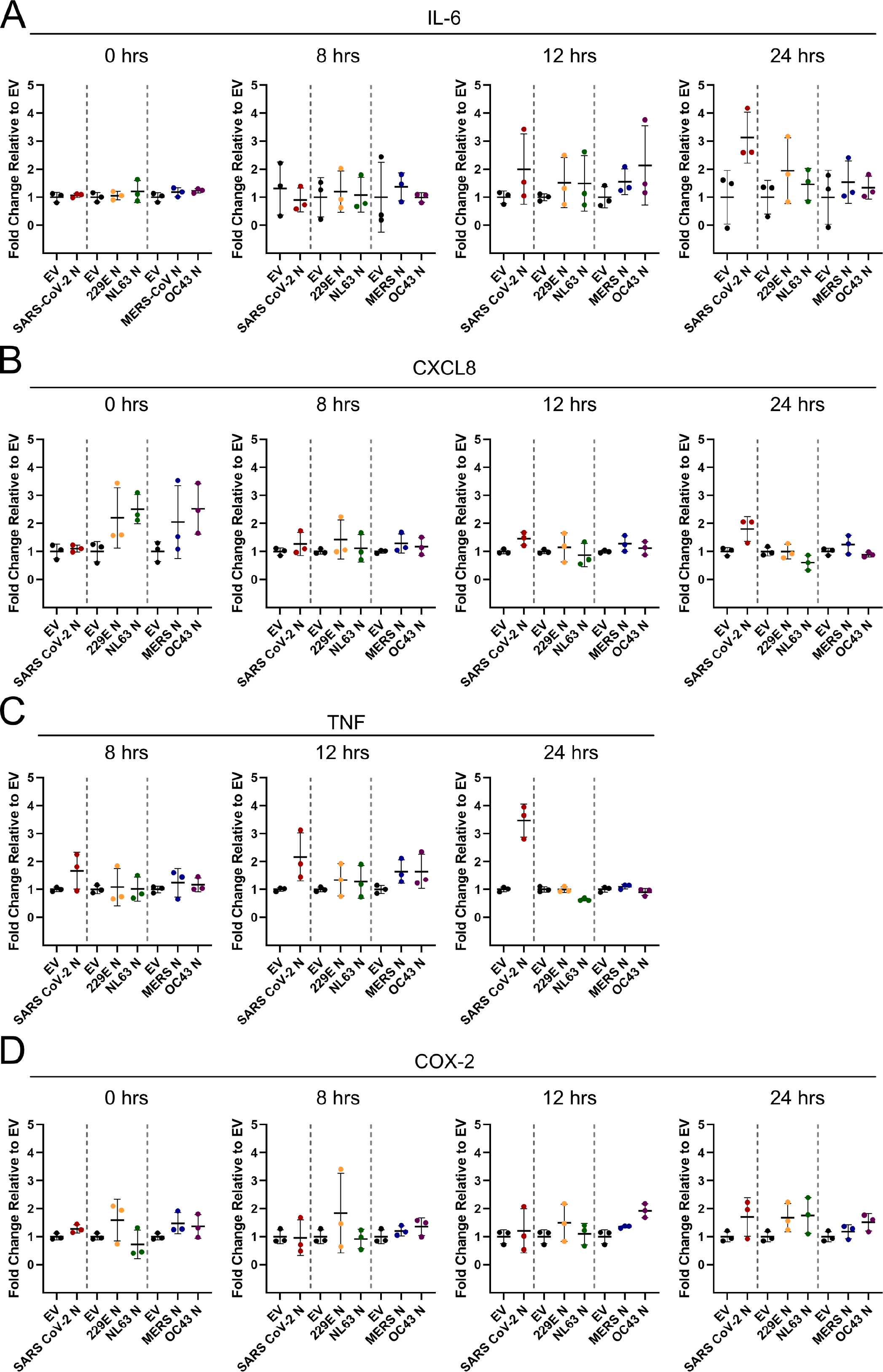
Ectopic expression of SARS-CoV-2 N elevates selected ARE-mRNAs. **A-C.** HUVECs were transduced with recombinant lentiviruses ectopically expressing alpha- and betacoronavirus N proteins or empty vector (EV) control, selected, and treated with 0.01 ng/L soluble TNF to increase transcription of ARE-containing cellular mRNAs. Total RNA was harvested at 0, 8, 12, and 24 hours post TNF treatment and RT-qPCR was performed using IL-6 (A), CXCL8 (B), TNF (C) and HPRT (house-keeping gene) specific primers. Values are represented as fold-change relative to EV-transduced cells for each time point. These data represent three independent experiments (*n=3*); mean ± SD.

Ectopic expression of SARS-CoV-2 N enhanced transcript levels of IL-6, CXCL8, and TNF 24 hours after transcription was induced; however, this enhancement was not deemed significant by statistical analysis (Fig 10A-C). In contrast, we did not observe any increase in transcript levels after expression of N protein from MERS-CoV, OC43, 229E, or NL63 (Fig 10A-C), consistent with earlier observations that their expression does not induce PB loss.

Taken together, we present the novel finding that three human CoVs induce PB loss and that for SARS-CoV-2, the nucleocapsid protein is responsible for this loss. Moreover, we tested five other human CoV N proteins and found that of these, only SARS-CoV-2 N was sufficient to induce significant PB disassembly and concomitantly redistributed and enhanced levels of two PB-localized cytokine mRNAs.

## Discussion

In this manuscript, we present the first evidence to show that human CoVs, including SARS-CoV-2, induce PB loss after infection. PBs are fundamental sites of post-transcriptional control of gene expression and are particularly relevant to the regulation of cytokine production. Our major findings are as follows: i) Three human coronaviruses, SARS-CoV-2, OC43, and 229E induced PB loss. ii) The SARS-CoV-2 nucleocapsid (N) protein was sufficient to cause PB loss. iii) N protein expression elevated levels of PB-localized cytokine transcripts encoding IL-6 and TNF. Taken together, these data point to PB loss as a feature of SARS-CoV-2 infection. Moreover, because viral induced PB disassembly elevates some PB-regulated cytokine transcripts, this phenotype may contribute to the dysregulation of proinflammatory molecules observed in severe SARS-CoV-2 infection.

We screened 27 SARS-CoV-2 gene products by transfection in HeLa cells [96] and initially identified eight candidate genes that reduced PB numbers (Fig 6). Validation of a subset of these in HUVECs revealed that the most robust and consistent mediator of PB loss was the SARS-CoV-2 viral N protein (Fig 7). The N protein is the most abundant produced protein during CoV replication [102]. The SARS-CoV-2 N protein is 419 amino acids long and has two globular and three intrinsically disordered protein domains, including a central disordered serine-arginine (SR-rich) linker region [102–104]. The N protein is a multifunctional RNA-binding protein (RBP) essential for viral replication; it coats the viral genome and promotes viral particle assembly [46,104–107]. Several recent reports have shown that N protein undergoes liquid-liquid phase separation with RNA, an event which may be regulated by phosphorylation of multiple serine residues in the SR-region and is an important feature of viral assembly [107–112] . The N protein is also an important modulator of antiviral responses [85,113,114]. A recent study showed that low doses of N protein supressed the production of IFN and some PB-regulated inflammatory cytokines, while high doses of N protein promoted their production [85] . These observations are consistent with our phenotype of PB disassembly, which correlates with later infection times, high expression of N protein and immunofluorescent staining throughout the cytoplasm (Fig 1, Fig 7).

We subsequently screened five other coronavirus N proteins from OC43, MERS, 229E, NL63, and SARS-CoV-1 discovered that the phenotype of N-mediated PB disassembly was not conserved among N proteins but was unique to SARS-CoV-2-N, and perhaps SARS-CoV-1 N protein which appeared to disassemble PBs in two of three independent experiments (Fig 8).

Despite conservation of structural motifs, the N proteins between most human CoVs possess low sequence conservation at the amino acid level (∼25-50%) and have been reported to exhibit different properties [115] . In contrast, SARS-CoV and SARS-CoV-2 N proteins have ∼94% amino acid identity [116], making it difficult to reconcile our observation that PB disassembly induced by SARS-CoV-2 N protein was significant, while PB disassembly induced by SARS-CoV-1 N protein was not (Fig 8). One difference that we observed by immunoblotting was the presence of a lower molecular weight ∼37 kDa band recognized by our anti-N antibody for SARS-CoV-2. We did not observe the 37kDa N product after OC43 infection, transduction with OC43 N protein or transduction with C-terminally Flag-tagged N proteins from 229E, NL63, MERS-CoV or SARS-CoV-1 (Fig 3, Fig 8). Steady state levels of the lower molecular weight N product increased over the course of SARS-CoV-2 infection, consistent with the timing of PB disassembly. Other groups have noted that the SARS-CoV-2 variant of concern (VOC), Alpha, produces an additional subgenomic mRNA from which a truncated version of N, termed N*, can be produced [68,117,118]. Translation of the N* ORF is predicted to start at an internal in-frame methionine residue (Met210) within the N protein [117, 118]. Alignment of the SARS-CoV-2 N protein sequence against other N proteins revealed that of the human CoVs, only SARS-CoV-1 and SARS-CoV-2 N retained a methionine at position 210 (229E: NC_002645.1, HKU1: NC_006577.2, MERS: NC_019843.3, NL63: NC_005831.2, OC43: NC_006213.1, SARS-1:

NC_004718.3, SARS-2: NC_045512.2). However, a lower molecular weight N protein product was not observed on our immunoblots for SARS-CoV N protein, leading us to speculate that the downstream Met residue may not be used for translation initiation in this case. Viruses capitalize on downstream methionine residues to translate truncated protein products with subcellular localization or functions that differ from their full-length counterparts as a clever way to increase coding capacity [119, 120] . Our ongoing investigation of the precise nature of the SARS-CoV-2 N protein truncation product we observe during infection and overexpression may reveal that it has a specific role in PB disassembly.

The PB protein MOV10, and other components of RNA processing machinery, were revealed as potential interactors with the N protein [121] ; however, we do not observe colocalization of N protein with PBs after immunofluorescent staining of SARS-CoV-2 infected cells or N-expressing cells. Based on our data, we consider two possible mechanisms of N protein mediated PB disassembly. First, N protein may mediate PB disassembly by phase separation with a PB protein(s). This is similar to what has already been shown for N-mediated disruption of cytoplasmic stress granules, important cytoplasmic biomolecular condensates that correlate with cellular translational shutdown [35]. N protein localizes to stress granules and binds the essential protein, G3BP1, preventing its interaction with other stress granule proteins and blocking stress granule formation [122–124]. Although the precise domain required for this effect has been debated, more than one report suggests that the N-terminal intrinsically disordered region is required for stress granule disruption [123, 125] . Second, a possible reason for PB loss may be the indiscriminate binding of RNA by N protein. N protein could be acting as sponge for RNA, pulling it out of cytoplasm, thereby reducing the RNA-protein interactions required for phase separation of PBs [24, 122]. We are currently engaged in site-directed and truncation mutagenesis studies to determine the precise region(s) of SARS-CoV-2-N that is essential for PB disassembly.

Prior to this report, little was known about CoVs and PBs, and the information that was published was contradictory. Infection with murine hepatitis virus (MHV) was reported to increase PBs, whereas transmissible gastroenteritis coronavirus (TGEV) decreased PBs [86, 87]. Since the initiation of our study, one additional publication used ectopic expression of one of the SARS-CoV-2 CoV proteases, nsp5, to test if it was capable of PB disassembly. Consistent with the results of our screen (Fig 6), nsp5 did not mediate PB loss [32]. In this manuscript, we now confirm that SARS-CoV-2, OC43, and 229E induce PB disassembly (Fig 1-3) [34] . We also observed that different human CoVs cause PB loss using different viral gene products; SARS-CoV-2 utilizes N protein but OC43 and 229E do not, a diversity which further underscores that convergence on PB disassembly by viruses likely benefits viral replication. Because PBs are composed of numerous cellular molecules with established (e.g. APOBEC, MOV10) or potential (decapping enzymes and exonucleases that could degrade viral RNA) antiviral activities, it is possible that viruses may target PBs for disassembly to negate their antiviral activity [28,29,31,39–45,126]. A recent screen for cellular proteins that bind SARS-CoV-2 viral RNA captured two PB proteins (Lsm14a and MOV10), which suggests CoV RNA may be shuttled to PBs [127]. That said, our evidence does not yet discern if the proposed antiviral role of PB-localized enzymes is promoted by phase separation of molecules into PBs or not; if so, we would predict that the antiviral function of these molecules is lost when PB granules decondense.

Emerging evidence suggests that activity of the decapping enzymatic complex is increased by phase separation and decreased in solution [9,24,128]. Thus, we speculate that PBs are direct-acting antiviral granules that can restrict virus infection when present as visible condensates; for this reason, they are targeted for disassembly by most viruses.

One possibility is that PBs are antiviral because their proteins help the cell respond to signals that activate innate immune pathways [28,30,129,130]. In support of this, TRAF6 was shown to control Dcp1a localization to PBs using ubiquitylation, suggesting that antiviral signaling is more complex than previously appreciated and integrates transcriptional responses with cytokine mRNA suppression in PBs [129, 131] . Moreover, the PB protein Lsm14A has been shown to bind to viral RNA/DNA after infection-induced PB disassembly to promote IRF3 activation and IFN-β production [130]. Although it remains unclear if the higher order condensation of many proteins into PBs is required for their proposed antiviral activity, what is clear is that the outcome of PB disassembly is a reversal of the constitutive decay or translational suppression of cytokine mRNA that would normally occur there [7,12–15,132] . Our data support a role for PBs as sensors of virus infection that release cytokine transcripts from repression when they disassemble. Using IF-FISH, we observed that RNAs encoding IL-6 and TNF, two molecules elevated in severe SARS-CoV-2 disease, localized to PBs. Upon SARS-CoV-2 N-induced PB disassembly, these transcripts re-localized from PB foci to the cytoplasm, a redistribution that was also accompanied by an increase in FISH probe signal intensity signifying increase RNA abundance (Fig 9). Although we did not see a statistically significant increase in IL-6 and TNF RNA level in SARS-CoV-2 N-expressing cells using a population-based assay like RT-qPCR (Fig 10), our single-cell analysis by IF-FISH suggests that the biological relevance of our observation derives from the combination of increased transcript level abundance and the redistribution of cytokine transcripts from PB foci to the cytoplasm. We speculate that viral infection causes PB loss and this event is viewed as a danger signal by the cell: it relieves cytokine mRNA suppression and re-localizes these transcripts, returning them to the cytoplasmic pool of RNA to permit translation of proinflammatory cytokines that then act as a call for reinforcements. In this way, PB disassembly is connected to the innate immune response and is one of many signals that notify the immune system that a cell is infected. In situations where interferon responses are delayed or defective, as is emerging for SARS-CoV-2 and severe COVID-19 [65–67], PB disassembly may occur to alert the immune system of an infection, and may be a contributing factor to pathogenic cytokine responses.

In summary, our work adds to a growing body of literature which describes that many viruses target PBs for disassembly, supporting the idea that PBs restrict viral infection. We showed that the N protein of SARS-CoV-2 is sufficient for PB disassembly, and this phenotype correlated with elevated levels of PB-localized cytokine transcripts encoding IL-6 and TNF, both which are elevated during SARS-CoV2 infection [53]. Not only does this work describe a previously uncharacterized cellular event that occurs during CoV infection, but we have identified a novel mechanism which may contribute to the dysregulated cytokine response exhibited by severe SARS-CoV-2 infection.

## Materials and Methods

### Cell Culture and Drug Treatments

All cells were maintained in humidified 37 °C incubators with 5% CO_2_ and 20% O_2_. Vero E6 (ATCC), HEK293T cells (ATCC), HeLa Tet-Off cells (Clontech) and HeLa Flp-In TREx GFP-Dcp1a cells (a generous gift from Dr. Anne-Claude Gingras) were cultured in DMEM (Thermo Fisher) supplemented with 100 U/mL penicillin, 100 µg/mL streptomycin, 2 mM L-glutamine (Thermo Fisher) and 10% FBS (Thermo Fisher). Calu3 (ATCC) and MRC-5 cells (ATCC, a generous gift from Dr. David Proud) were cultured in EMEM (ATCC) supplemented with 100 U/mL penicillin, 100 µg/mL streptomycin, 2 mM L-glutamine and 10% FBS (Thermo Fisher). HUVECs (Lonza) were cultured in endothelial cell growth medium (EGM-2) (Lonza). HUVECs were seeded onto tissue culture plates or glass coverslips coated with 0.1 % (w/v) porcine gelatin (Sigma) in 1x PBS. For sodium arsenite treatment, HUVECs were treated with 0.25 mM sodium arsenite (Sigma) or a vehicle control for 30 min.

### Plasmids and Cloning

pLenti-IRES-Puro SARS-CoV-2 plasmids were a generous gift from the Krogan Lab [96]. pLJM1-OC43-N was cloned from pGBW-m4134906, a gift from Ginkgo Bioworks & Benjie Chen (Addgene plasmid #151960; http://n2t.net/addgene:151960; RRID:Addgene_151960) using BamHI and EcoRI restriction sites (NEB). pLJM1-NL63-N-FLAG was cloned from pGBW-m4134910, a gift from Ginkgo Bioworks & Benjie Chen (Addgene plasmid #151939; http://n2t.net/addgene:151939; RRID:Addgene_151939) using BamHI and EcoRI. pLJM1-229E-N-FLAG was cloned from pGBW-m4134902, a gift from Ginkgo Bioworks & Benjie Chen (Addgene plasmid #151912; http://n2t.net/addgene:151912; RRID:Addgene151912) using BamHI and EcoRI. Codon-optimized pLJM1-MERS-CoV-N-FLAG was cloned from SinoBiological (cat #VG40068-CF) using BamHI and EcoRI (Table 1). Codon-optimized pLJM1-SARS-CoV-N-FLAG was cloned SinoBiological (cat# VG40588-NF) using BamHI and EcoRI. The Kaposi’s sarcoma-associated herpesvirus Kaposin B (KapB, KS lung isolate) clone in pLJM1 was previously described [91].

**Table 1:** Plasmids.

| Plasmid | Use | Source | Mammalian Selection |
| --- | --- | --- | --- |
| pLJM1-Puro | Empty Vector Control | [90] [129] | Puromycin |
| pLJM1-BSD | Empty Vector Control | [129] | Blasticidin |
| pLJM1-ACE2 | Overexpression |  | Blasticidin |
| pMD2.G | Lentivirus Generation | Addgene #12259 | N/A |
| psPAX2 | Lentivirus Generation | Addgene #12260 | N/A |
| pLJM1-OC43-N | Overexpression | Addgene #151960 | Puromycin |
| pLJM1-229E-N-FLAG | Overexpression | Addgene #151912 | Puromycin |
| pLJM1-NL63-N-FLAG | Overexpression | Addgene #151939 | Puromycin |
| pLJM1-MERS-CoV-N-FLAG | Overexpression | SinoBiological<br>#VG40068-CF | Puromycin |
| pLJM1-SARS-CoV-1-N-FLAG | Overexpression | SinoBiological<br># VG40588-NF | Puromycin |
| pLenti-SARS-CoV-2-IRES-strep | Overexpression library | N Krogan (UCSF)<br>[114] | Puromycin |

### Transient Transfections

Transient transfections were performed using Fugene (Promega) according to manufacturer’s guidelines. Briefly, HeLa Flp-In TREx GFP-Dcp1a cells were seeded in 12-well plates at 150,000 cells/well in antibiotic-free DMEM. Cells were transfected with 1 μg of DNA and 3 μL of Fugene for 48 hours before processing.

### Production and use of Recombinant Lentiviruses

All recombinant lentiviruses were generated using a second-generation system. HEK293T cells were transfected with psPAX2, MD2-G, and the lentiviral transfer plasmid containing a gene of interest using polyethylenimine (PEI, Polysciences). 6 hours after transfection, serum-free media was replaced with DMEM containing serum but no antibiotics. Viral supernatants were harvested 48 hours post-transfection and frozen at -80°C until use. For transduction, lentiviruses were thawed at 37°C and added to target cells in complete media containing 5 µg/mL polybrene (Sigma). After 24 hours, the media was replaced with selection media containing 1 µg/mL puromycin or 5 µg/mL blasticidin (ThermoFisher) and cells were selected for 48 h before proceeding with experiments.

### Immunofluorescence

Cells were seeded onto 18mm round, #1.5 coverslips (Electron Microscopy Sciences) for immunofluorescence experiments. Following treatment, cells were fixed for 10 or 30 (if infected with SARS-CoV-2) min in 4% (v/v) paraformaldehyde (Electron Microscopy Sciences).

Samples were permeabilized with 0.1% (v/v) Triton X-100 (Sigma-Aldrich) for 10 min at room temperature and blocked in 1% human AB serum (Sigma-Aldrich) in 1X PBS 1 h at room temperature. Primary and secondary antibodies (Table 2) were diluted in 1% human AB serum and used at the concentrations in Table 2. Nuclei were stained with 1 µg/ml Hoechst (Invitrogen). Samples were mounted with Prolong Gold AntiFade mounting media (ThermoFisher). Images were captured using a Zeiss AxioObserver Z1 microscope with the 40X oil-emersion objective unless otherwise stated in the respective figure caption. To account for variability in staining, all experiments contained internal controls (negative control; mock or EV). During image acquisition, exposure time was kept consistent within an independent experiment.

**Table 2:**
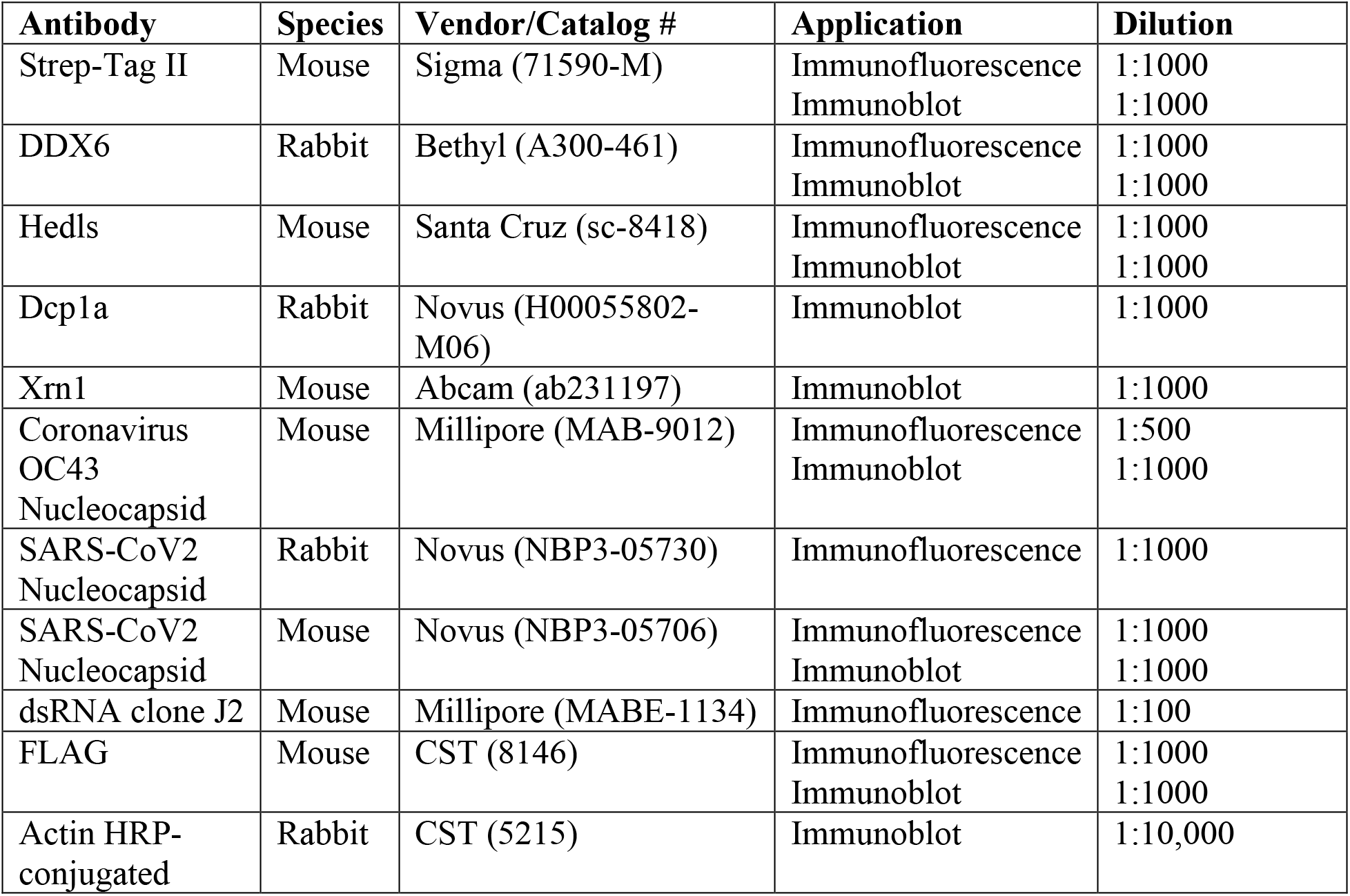
Antibodies.

### Immunofluorescence (IF) RNA Fluorescent *In Situ* Hybridization (FISH)

IF-FISH was performed according to manufacturers protocol (Stellaris IF-FISH protocol). Briefly, cells were fixed in 4% formaldehyde (Sigma-Aldrich) for 10 mins then permeabilized with 0.1% Triton X-100 (Sigma-Aldrich) in PBS for 10 mins. Primary Hedls antibody was diluted in PBS and incubated for 1 h at room temp, following which secondary antibody was likewise diluted in PBS and allowed to incubate for 1 h at room temp. Cells were again fixed in 4% formaldehyde for 10 mins, washed with 1X PBS, and incubated in Wash Buffer A (LGC Biosearch) for 5 minutes. Stellaris probes; GAPDH Quasar 670 (VSMF 2151-5), IL-6 Quasar 670 (LGC Biosearch, SMF 1065-5, custom order, probe sequences detailed in Table 4) and TNF Quasar-670 (LGC Biosearch, SMF 1065-5, custom order, probe sequences detailed in Table 4). were diluted in Hybridization Buffer (LGC Biocearch) to a concentration of 125 nM and cells were incubated for hybridization at 37 °C overnight. The following day, cells were washed sequentially with Wash Buffer A for 30 mins, Wash Buffer A containing 5 ng/mL DAPI (Invitrogen) for 30 mins, and Wash Buffer B (LGC Biosearch) for 5 minutes. Samples were mounted with Prolong Gold AntiFade mounting media (ThermoFisher). Images were captured on a Zeiss LSM 880 confocal microscope with the 63X oil-emersion objective. At least three z-stacks were acquired per condition, maximal intensity projections (MIPs) are presented.

### Immunoblotting

Cells were lysed in 2X Laemmli buffer and stored at -20°C until use. The DC Protein Assay (Bio-Rad) was used to quantify protein concentration as per the manufacturer’s instructions. 10-15 µg of protein lysate was resolved by SDS-PAGE on TGX Stain-Free acrylamide gels (BioRad). Total protein images were acquired from the PVDF membranes after transfer on the ChemiDoc Touch Imaging system (BioRad). Membranes were blocked in 5% BSA in TBS-T (Tris-buffered saline 0.1% Tween-20). Primary and secondary antibodies were diluted in 2.5% BSA, dilutions can be found in Table 2. Membranes were visualized using Clarity Western ECL substrate and the ChemiDoc Touch Imaging system (BioRad). Densitometry was conducted using ImageJ, area under the curve for each band was calculated, normalized to the respective actin loading control band, and presented as fold change.

### Quantitative PCR

RNA was collected using the RNeasy Plus Mini Kit (Qiagen) according to the manufacturer’s instructions and stored at -80°C until further use. RNA concentration was determined using NanoDrop One^C^ (ThermoFisher) and 500 ng of RNA was reverse transcribed using qScript XLT cDNA SuperMix (QuantaBio) using a combination of random hexamer and oligo dT primers, according to the manufacturer’s instructions. Depending on starting concentration, cDNA was diluted between 1:10 and 1:20 for qPCR experiments and SsoFast EvaGreen Mastermix (Biorad) was used to amplify cDNA. The ΔΔquantitation cycle (Cq) method was used to determine the fold change in expression of target transcripts using HPRT as a housekeeping control gene. Variance in the mock or empty vector samples was calculated by dividing the ΔCt value of a single replicate by the average ΔCt value of all replicates for that specific gene and condition. qPCR primer sequences can be found in Table 3. (Table 3).

**Table 3:**
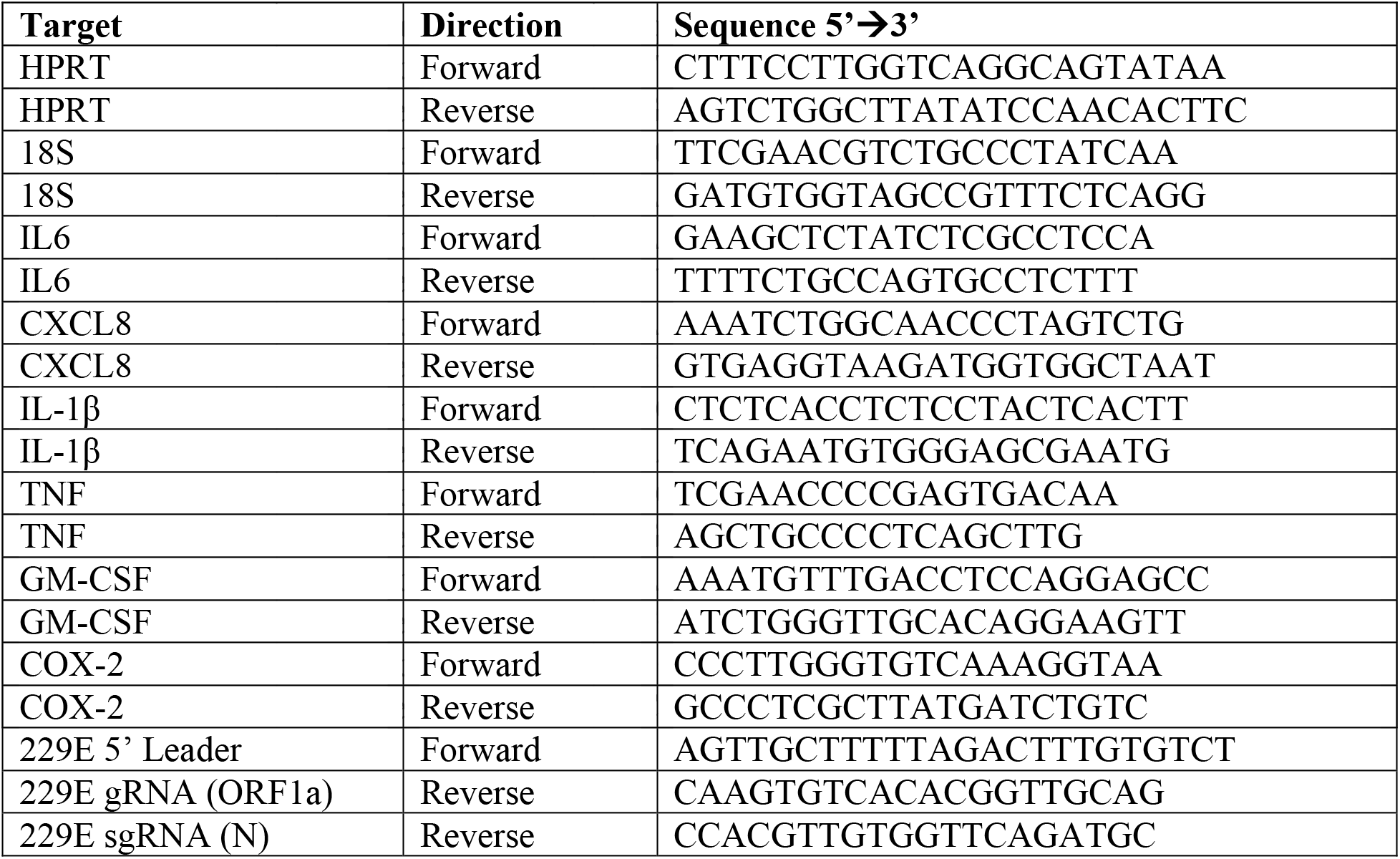
qPCR primers.

**Table 4.**
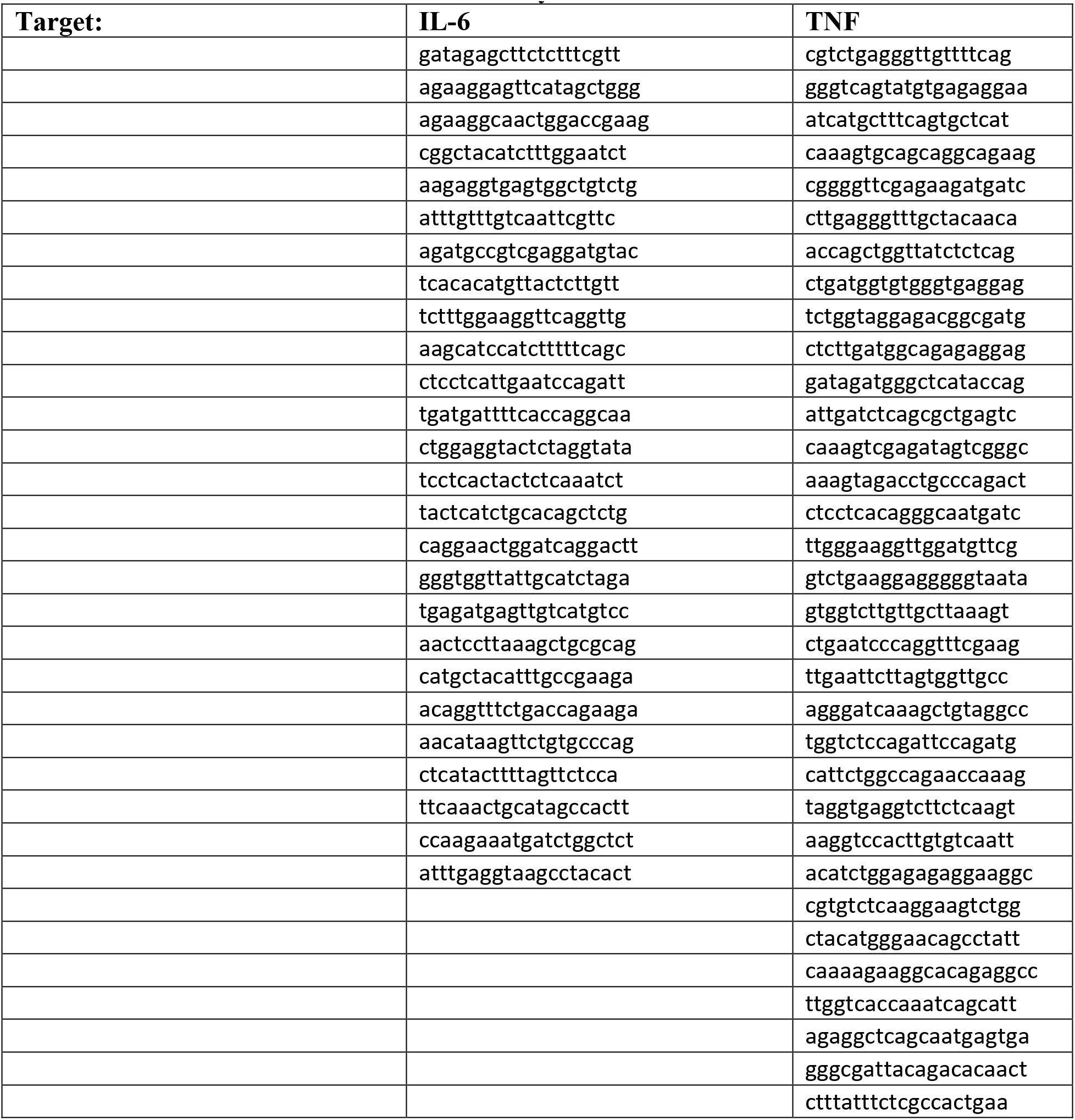
Probes for RNA Fluorescent in situ hybridization.

### Virus Stocks and Virus Propagation

Experiments with SARS-CoV-2 and variants were conducted in a containment level-3 (CL3) facility, and all standard operating procedures were approved by the CL3 Oversight Committee and Biosafety Office at the University of Calgary. Stocks of SARS-CoV-2 Toronto-01 isolate (SARS-CoV-2/SB3-TYAGNC) [90], Alpha, Beta, Gamma and Delta were propagated in Vero E6 cells. To produce viral stocks, Vero E6 cells were infected at an MOI of 0.01 for 1 hour in serum-free DMEM at 37°C. Following adsorption, DMEM supplemented with 2% heat inactivated FBS and 100 U/mL penicillin/streptomycin/glutamine was added to the infected wells. 24-60 hours post-infection (hpi), the supernatant was harvested and centrifuged at 500 x *g* for 5 min to remove cellular debris. Virus stocks were aliquoted and stored at -80°C for single use. SARS-CoV-2 titres were enumerated using plaque assays on Vero E6 cells as previously described [133] using equal parts 2.4% w/v semi-solid colloidal cellulose overlay (Sigma; prepared in ddH_2_O) and 2X DMEM (Wisent) with 1% heat inactivated FBS and 1% PSQ. Experiments with hCoV-OC43 (ATCC VR-1558) and hCoV-229E (ATCC VR-740) were conducted in under containment level-2 conditions. hCoV-OC43 and hCoV-229E were propagated in Vero E6 and MRC-5 cells, respectively. Cells were infected at an MOI of 0.01 for 1 hour in serum-free media at 33°C. Following adsorption, the viral inoculum was removed and replaced with fresh media supplemented with 2% heat inactivated FBS and 100 U/mL penicillin/streptomycin/glutamine. After 5-6 days post infection (dpi), the supernatant was harvested and cellular debris was cleared by centrifugation. Virus stocks were aliquoted and stored at -80°C. hCoV-OC43 and hCoV-229E titres were enumerated using Reed and Muench tissue-culture infectious dose 50% (TCID_50_) in Vero E6 or MRC-5 cells, respectively.

### Virus Infection

For experimental infections, cells were seeded into wells to achieve ∼80% confluency at the time of infection. The growth media was removed and replaced with 100 µL of viral inoculum diluted in serum-free DMEM to reach the desired MOI and incubated at 37°C for 1 hour, rocking the plate every 10 min. Following incubation, the virus inoculum was removed and replaced with 1 mL of complete growth media.

### Processing Body and FISH Probe Intensity Quantification

Processing bodies were quantified using an unbiased image analysis pipeline generated in the freeware CellProfiler4.0.6 (cellprofiler.org) [134]. First, nuclear staining was used to identify individual cells applying a binary threshold and executing primary object detection between 65 and 200 pixels for HUVECs/HUVEC^ACE2^ and 40 and 200 pixels for Calu3s. For each identified object (nuclei), the peripheral boundary of each cell was defined using the “Propagation” function. Propagation distance was customized depending on cell type to account for variance in cell size (150 pixel radius from the nuclei for HUVECs/HUVEC^ACE2^, 60 pixel radius from nuclei for Calu3s). Using a subtractive function to remove the nuclei from the total cell area, the cytoplasm of each cell was defined. The cytoplasm area mask was then applied to the matched image stained for PB proteins (DDX6 or Hedls) to count only cytoplasmic puncta. Importantly, multiple nuclei in close proximity (e.g., within the propagation distance) would be divided into mutually exclusive cells, such that a single PB could not be counted more than once. Cells were stratified into ‘positive cells’ (staining for viral proteins or dsRNA) or ‘bystander cells’. In control treatments (e.g., mock infected or EV) PBs in all cells were quantified. In treatment cells only ‘positive cells’ were quantified unless otherwise indicated in corresponding figure captions. Background staining was reduced using the “Enhance Speckles” function. Only DDX6 or Hedls-positive puncta with a defined size (3-13 pixels) and intensity range were quantified using “global thresholding with robust background adjustments” function. All thresholds were consistent within each replicate that used identical staining parameters. PBs were quantified per cell, either control (EV or mock) or treatment (infected or viral protein expressing). For each experiment, an equal number of control and treatment cells were analyzed. For Figure 5, puncta counts were exported and RStudio was used for data analysis and PB numbers in treatment cells were represented relative to control cells for ease of data interpretation, rather than raw PBs per cell. For Figure 9, nuclei and cells were defined according to HUVEC boundaries as specified above. Background of probe specific staining was reduced using a global thresholding strategy with a minimum cross entropy method of execution. The MeasureImageIntensity function was then used to identify the total intensity per image which was divided by the number of cells identified within that image for a final readout of total intensity/cell. A minimum of 40 cells were quantified per condition.

### Statistics

All statistical analyses were performed using GraphPad Prism 9.0. ‘Per cell’ processing body counts were plotted such that each independent biological replicate (including paired control and treatment) could be visualized on a single graph. Given that per cell processing body counts are naturally skewed and thus non-parametric, we elected to use rank-sum statistical methods (Mann-Whitney U test and Kruskal Wallis test) when appropriate, as indicated in the corresponding figure caption. In the case of Figure 7, a two-way ANOVA with a Tukey’s post-hoc analysis was used to determine significance to (1) avoid the false discovery rate associated with multiple T-tests and (2) because sodium arsenite-induced processing bodies appeared to be parametrically distributed. Parametric distribution was assumed on all normalized data (Fig 5, 6, 9, 10, S4, & S7) and therefore we elected to use unpaired T-tests or one-way ANOVAs with Dunnett’s post-hoc analysis, as indicated in the corresponding figure caption.

## Supplemental Figure Legends

**Figure S1. HUVEC-ACE2 cells are permissive to SARS-CoV-2. A.** HUVECs were transduced with recombinant lentiviruses expressing human ACE2, selected, and infected with SARS-CoV-2 (TO-1 isolate; MOI=3). 6- and 24-hours post-infection, virus-containing supernatant was harvested and titrated by TCID_50_ assay in VeroE6 cells. These data represent two independent experiments (*n*=2); mean ± SD. **B-C.** HUVECs were transduced with ACE2 as in A. Cells were fixed and immunostained for DDX6 (B) or Hedls (C) and quantified per field of view using CellProfiler. These data represent two independent experiments (*n*=2) with >100 cells measured per condition. Each EV and ACE2 replicate pair plotted independently; mean. Statistics were performed using a Mann-Whitney rank-sum test (ns, nonsignificant).

**Figure S2. HUVECs are permissive to OC43 and 229E infection. A-B.** HUVECs were infected with OC43 (TCID_50_ = 2 x 10^4^) or 229E (TCID_50_ = 2.4 x 10^3^). 6-, 12-, or 24 hpi, virus-containing supernatant was harvested and titrated TCID_50_ assay in VeroE6 (A) or MRC5 (B) cells, respectively. These data represent two independent experiments (*n*=2); mean ± SD. **C.** HUVECs were infected with 229E as in B or mock infected. Total RNA was harvested 12 hpi and RT-qPCR was performed using primers specific to 229E genomic RNA (gRNA) or subgenomic RNA8 (sgRNA8), or HPRT (cellular house-keeping gene). Cycle-threshold (Ct) values for each primer pair were plotted. These data represent three independent experiments (*n=*3); mean ± SD. Ct values greater than 35 were not deemed biologically relevant and therefore are scored as N/A. **D.** PCR products from C. were resolved using agarose gel electrophoresis. gRNA fragment migrated at the expected size of 285 bp. sgRNA fragment migrated at its expected ∼80 bp. Representative images from one of two independent experiments are shown.

**Figure S3. Hedls puncta are absent in OC43 and 229E infected cells. A.** HUVECs were infected with OC43 (TCID_50_ = 2 x 10^4^) or mock-infected. 12 or 24 hpi cells were fixed and immunostained for DDX6 (green) and Hedls (white). Nuclei were stained with Hoechst (blue). Representative images from one of three independent experiments shown. **B.** Hedls puncta in OC43-infected or mock-infected cells were quantified per field of view using CellProfiler. **C.** HUVECs were infected with 229E (TCID_50_ = 2.4 x 10^3^) or mock-infected. 6 or 12 hpi cells were fixed and immunostained for DDX6 (green) and Hedls (white). Nuclei were stained with Hoechst (blue). Representative images from one of three independent experiments shown. **D.** Hedls puncta in 229E-infected or mock-infected cells were quantified as in B. These data represent three independent experiments (*n=*3) with ≥100 cells measured per condition per replicate. Each mock and infected replicate pair plotted independently; mean. Statistics were performed using a Mann-Whitney rank-sum test (****, *p* < 0.0001). Scale bar = 20 μm.

**Figure S4. Processing body protein densitometry after human coronavirus infection. A.** HUVECs were transduced with human ACE2, selected, and infected with SARS-CoV-2 TO-1 isolate (MOI=3). Cells were lysed at 6 and 12 hpi and immunoblotting was performed using XRN1, Hedls, DCP1A, DDX6, SARS-CoV-2 N, and β-actin specific antibodies. The results of second independent experiment of two is shown. **B-C.** HUVECs were infected with OC43 (TCID_50_ = 2 x 10^4^) or 229E (TCID_50_ = 2.4 x 10^3^) or mock-infected. Cells were lysed at 12 and 24 hpi (B, OC43) or 6 and 12 hpi (C, 229E). Immunoblotting was performed using DDX6, Hedls, XRN1, DCP1A, and actin. Protein densitometry was determined in ImageJ normalized to actin and expressed as a fold-change relative to the 12 hpi (B; OC43) or 6 hpi (C; 229E) mock-infected control. These data represent three independent biological replicates (*n*=3). A one-way ANOVA with a Dunnett’s post-hoc analysis was performed; mean; bars represent SD (*, *p* < 0.05).

**Figure S5. Ectopic expression of SARS-CoV-2 ORFs**. HeLa cells expressing GFP-Dcp1a were transfected with an empty vector (EV) control or 2xStrep-tagged SARS-CoV-2 ORFs for 48 hours then fixed and immunostained for Strep-Tag-II (viral ORF; green) or DDX6 (PBs; white). Nuclei were stained with Hoechst (blue). A. SARS-CoV-2 ORFs that could be detected by Strep-Tag-II staining (thresholded ORFs). B. SARS-CoV-2 ORFs that could not be detected by Strep-Tag-II staining (unthresholded ORFs). Representative images from one of three or more independent experiments shown. Scale is the same for all images. N=nucleocapsid protein, E=envelope protein, M=membrane protein, S=spike protein

**Figure S6. Validation of processing body loss induced by selected SARS-CoV-2 ORFs in HUVECs. A.** HUVECs were transduced with recombinant lentiviruses expressing 2xStrep-tagged SARS-CoV-2 nsp1, nsp14, ORF3b, and ORF7b constructs or control lentiviruses (EV) and selected with puromycin for 48 hours. Cells were fixed and immunostained for Strep-Tag-II (ORFs; green) and DDX6 (PBs; white). Nuclei were stained with Hoechst (blue). Representative images from one of three independent experiments shown. Scale bar = 20 µm. **B.** DDX6 puncta were quantified per field of view using CellProfiler. These data represent three independent experiments (*n=*3) with >18 cells measured per condition per replicate. Each EV and ORF-expressing replicate pair plotted independently; mean. Statistics were performed using Kruskal-Wallis H test with Dunn’s correction (**, *p* < 0.0021; ****, *p* < 0.0001; ns, nonsignificant). **C.** HUVECs were transduced as in A. Cells were lysed and immunoblotting was performed using the Strep-Tag II antibody on a 4-15% gradient gel.

**Figure S7. Processing body protein densitometry after CoV N protein overexpression. A-C.** HUVECs were transduced with recombinant lentiviruses ectopically expressing N protein from the betacoronaviruses MERS-CoV and OC43 (A), N protein from alphacoronaviruses 229E and NL63 (B) or SARS-CoV-1 N (C). A control lentiviral expressing an empty vector (EV) was used as a negative control and SARS-CoV-2 N protein expressing lentiviruses were used as a positive control in each experiment. Cells were selected and lysed and immunoblotting was performed using XRN1, Hedls, DCP1A, DDX6, N protein or FLAG, and β-actin specific antibodies. Protein densitometry was determined in ImageJ normalized to actin and expressed as a fold-change relative to the EV control. These data represent three independent biological replicates (*n*=3). A one-way ANOVA with a Dunnett’s post-hoc analysis was performed; mean; bars represent SD (*, *p* < 0.05).

**Figure S8. TNF treatment does not alter processing body number**

HUVECs were treated with 0.01 ng/L soluble TNF for either 6 or 12 hours following which, cells were fixed and immunostained for Hedls. Nuclei were stained with Hoechst. Hedls puncta were quantified per field of view using CellProfiler. These data represent two independent experiments (*n=*2) with >90 cells measured per condition per replicate. Each mock and infected replicate pair plotted independently; mean.

## Acknowledgements

We sincerely thank the members of the Corcoran lab, Dr. Craig McCormick and Dr. Eric Pringle for helpful discussions about this work. We would like to thank Dr. Nevan Krogan (UCSF) for the SARS-CoV-2 gene library, Dr. David Proud (University of Calgary) for MRC-5 cells, and Dr. Lorne Tyrrell, Dr. Michael Joyce and Holly Bandi (University of Alberta) for isolates of SARS-CoV-2 variants Alpha, Beta, Gamma and Delta. We would like to thank Dr. Anne Vaahtokari of the Charbonneau Microscopy Facility for microscopy support and Dr. Devender Kumar for CL3 facility support. MK was supported by a CSM graduate training award, a CIHR CGS-M scholarship, and a CIHR doctoral award and RPM was supported by a Snyder Institute Beverley Phillips Doctoral training award. This study was supported by operating funds awarded to JAC from the Canadian Institutes for Health Research (CIHR): a COVID rapid response operating grant (#177704) and an operating grant (#175622) the Coronavirus Variants Rapid Response Network (CoVaRR-Net), of which JAC is a member. The Vaccine and Infectious Disease Organization (VIDO) receives operational funding for its CL3 facility (InterVac) from the Canada Foundation for Innovation through the Major Science Initiatives. VIDO also receives operational funding from the Government of Saskatchewan through Innovation Saskatchewan and the Ministry of Agriculture.

## Author Contributions

Mariel Kleer: Conceptualization, Experimentation, Analysis, Paper Writing

Rory P. Mulloy: Conceptualization, Experimentation, Analysis, Paper Writing

Carolyn-Ann Robinson: Conceptualization, Experimentation, Analysis, Paper Writing

Danyel Evseev: Experimentation, Analysis, Paper Writing

Maxwell P. Bui-Marinos: Experimentation, Analysis, Paper Writing

Elizabeth L. Castle: Analysis

Arinjay Banerjee: Reagents, Paper Editing

Samira Mubareka: Reagents

Karen Mossman: Reagents

Jennifer A. Corcoran: Conceptualization, Experimentation, Supervision, Funding Acquisition,

Project Administration, Paper Writing

## References

1. Eulalio A, Behm-Ansmant I, Schweizer D, Izaurralde E. P-body formation is a consequence, not the cause, of RNA-mediated gene silencing. Mol Cell Biol. 2007;27: 3970–3981. doi:10.1128/MCB.00128-07

2. Eulalio A, Rehwinkel J, Stricker M, Huntzinger E, Yang S-F, Doerks T, et al. Target-specific requirements for enhancers of decapping in miRNA-mediated gene silencing. Genes Dev. Cold Spring Harbor Lab; 2007;21: 2558–2570. doi:10.1101/gad.443107

3. Hubstenberger A, Courel M, Bénard M, Souquere S, Ernoult-Lange M, Chouaib R, et al. P-Body Purification Reveals the Condensation of Repressed mRNA Regulons. Mol Cell. Elsevier Inc; 2017;68: 144–157.e5. doi:10.1016/j.molcel.2017.09.003

4. Youn J-Y, Dunham WH, Hong SJ, Knight JDR, Bashkurov M, Chen GI, et al. High-Density Proximity Mapping Reveals the Subcellular Organization of mRNA-Associated Granules and Bodies. Mol Cell. Elsevier Inc; 2018;69: 517–532.e11. doi:10.1016/j.molcel.2017.12.020

5. Bakheet T, Hitti E, Khabar KSA. ARED-Plus: an updated and expanded database of AU-rich element-containing mRNAs and pre-mRNAs. Nucleic Acids Research. 2018;46: D218–D220. doi:10.1093/nar/gkx975

6. Hitti E, Bakheet T, Al-Souhibani N, Moghrabi W, Al-Yahya S, Al-Ghamdi M, et al. Systematic Analysis of AU-Rich Element Expression in Cancer Reveals Common Functional Clusters Regulated by Key RNA-Binding Proteins. Cancer Res. 2016;76: 4068–4080. doi:10.1158/0008-5472.CAN-15-3110

7. Blanco FF, Sanduja S, Deane NG, Blackshear PJ, Dixon DA. Transforming growth factor β regulates P-body formation through induction of the mRNA decay factor tristetraprolin. Mol Cell Biol. 2014;34: 180–195. doi:10.1128/MCB.01020-13

8. Brengues M, Teixeira D, Parker R. Movement of eukaryotic mRNAs between polysomes and cytoplasmic processing bodies. Science. American Association for the Advancement of Science; 2005;310: 486–489. doi:10.1126/science.1115791

9. Riggs CL, Kedersha N, Ivanov P, Anderson P. Mammalian stress granules and P bodies at a glance. J Cell Sci. 2020;133: jcs242487. doi:10.1242/jcs.242487

10. Khong A, Parker R. mRNP architecture in translating and stress conditions reveals an ordered pathway of mRNP compaction. J Cell Biol. 2018;217: 4124–4140. doi:10.1083/jcb.201806183

11. Matheny T, Rao BS, Parker R. Transcriptome-Wide Comparison of Stress Granules and P-Bodies Reveals that Translation Plays a Major Role in RNA Partitioning. Mol Cell Biol. 2019;39: 612. doi:10.1128/MCB.00313-19

12. Corcoran JA, Khaperskyy DA, Johnston BP, King CA, Cyr DP, Olsthoorn AV, et al. Kaposi’s sarcoma-associated herpesvirus G-protein-coupled receptor prevents AU-rich-element-mediated mRNA decay. J Virol. 2012;86: 8859–8871. doi:10.1128/JVI.00597-12

13. Corcoran JA, Johnston BP, McCormick C. Viral Activation of MK2-hsp27-p115RhoGEF-RhoA Signaling Axis Causes Cytoskeletal Rearrangements, P-body Disruption and ARE-mRNA Stabilization. Robertson ES, editor. PLoS Pathogens. Public Library of Science; 2015;11: e1004597. doi:10.1371/journal.ppat.1004597

14. Franks TM, Lykke-Andersen J. TTP and BRF proteins nucleate processing body formation to silence mRNAs with AU-rich elements. Genes Dev. 2007;21: 719–735. doi:10.1101/gad.1494707

15. Vindry C, Marnef A, Broomhead H, Twyffels L, Ozgur S, Stoecklin G, et al. Dual RNA Processing Roles of Pat1b via Cytoplasmic Lsm1-7 and Nuclear Lsm2-8 Complexes. CellReports. 2017;20: 1187–1200. doi:10.1016/j.celrep.2017.06.091

16. Vandelli A, Cid Samper F, Torrent Burgas M, Sanchez de Groot N, Tartaglia GG. The Interplay Between Disordered Regions in RNAs and Proteins Modulates Interactions Within Stress Granules and Processing Bodies. J Mol Biol. 2021;: 167159. doi:10.1016/j.jmb.2021.167159

17. Nosella ML, Forman-Kay JD. Phosphorylation-dependent regulation of messenger RNA transcription, processing and translation within biomolecular condensates. Curr Opin Cell Biol. 2021;69: 30–40. doi:10.1016/j.ceb.2020.12.007

18. Jalihal AP, Pitchiaya S, Xiao L, Bawa P, Jiang X, Bedi K, et al. Multivalent Proteins Rapidly and Reversibly Phase-Separate upon Osmotic Cell Volume Change. Mol Cell. 2020;79: 978–990.e5. doi:10.1016/j.molcel.2020.08.004

19. Mitrea DM, Chandra B, Ferrolino MC, Gibbs EB, Tolbert M, White MR, et al. Methods for Physical Characterization of Phase-Separated Bodies and Membrane-less Organelles. J Mol Biol. 2018;430: 4773–4805. doi:10.1016/j.jmb.2018.07.006

20. Banani SF, Rice AM, Peeples WB, Lin Y, Jain S, Parker R, et al. Compositional Control of Phase-Separated Cellular Bodies. Cell. 2016;166: 651–663. doi:10.1016/j.cell.2016.06.010

21. Guillén-Boixet J, Kopach A, Holehouse AS, Wittmann S, Jahnel M, Schlüßler R, et al. RNA-Induced Conformational Switching and Clustering of G3BP Drive Stress Granule Assembly by Condensation. Cell. 2020;181: 346–361.e17. doi:10.1016/j.cell.2020.03.049

22. Van Treeck B, Parker R. Emerging Roles for Intermolecular RNA-RNA Interactions in RNP Assemblies. Cell. 2018;174: 791–802. doi:10.1016/j.cell.2018.07.023

23. Cougot N, Cavalier A, Thomas D, Gillet R. The dual organization of P-bodies revealed by immunoelectron microscopy and electron tomography. J Mol Biol. 2012;420: 17–28. doi:10.1016/j.jmb.2012.03.027

24. Corbet GA, Parker R. RNP Granule Formation: Lessons from P-Bodies and Stress Granules. Cold Spring Harb Symp Quant Biol. 2019;84: 203–215. doi:10.1101/sqb.2019.84.040329

25. Standart N, Weil D. P-Bodies: Cytosolic Droplets for Coordinated mRNA Storage. Trends Genet. 2018;34: 612–626. doi:10.1016/j.tig.2018.05.005

26. Docena G, Rovedatti L, Kruidenier L, Fanning A, Leakey NAB, Knowles CH, et al. Down-regulation of p38 mitogen-activated protein kinase activation and proinflammatory cytokine production by mitogen-activated protein kinase inhibitors in inflammatory bowel disease. Clin Exp Immunol. John Wiley & Sons, Ltd; 2010;162: 108–115. doi:10.1111/j.1365-2249.2010.04203.x

27. Jangra RK, Yi M, Lemon SM. DDX6 (Rck/p54) is required for efficient hepatitis C virus replication but not for internal ribosome entry site-directed translation. J Virol. 2010;84: 6810–6824. doi:10.1128/JVI.00397-10

28. Ng CS, Kasumba DM, Fujita T, Luo H. Spatio-temporal characterization of the antiviral activity of the XRN1-DCP1/2 aggregation against cytoplasmic RNA viruses to prevent cell death. Cell Death & Differentiation. Nature Publishing Group; 2020;36: 932–20. doi:10.1038/s41418-020-0509-0

29. Ostareck DH, Naarmann-de Vries IS, Ostareck-Lederer A. DDX6 and its orthologs as modulators of cellular and viral RNA expression. Wiley Interdiscip Rev RNA. 2014;5: 659–678. doi:10.1002/wrna.1237

30. Dougherty JD, Tsai W-C, Lloyd RE. Multiple Poliovirus Proteins Repress Cytoplasmic RNA Granules. Viruses. Multidisciplinary Digital Publishing Institute; 2015;7: 6127– 6140. doi:10.3390/v7122922

31. Dougherty JD, White JP, Lloyd RE. Poliovirus-mediated disruption of cytoplasmic processing bodies. J Virol. 2011;85: 64–75. doi:10.1128/JVI.01657-10

32. Fan S, Xu Z, Liu P, Qin Y, Chen M. Enterovirus 71 2A Protease Inhibits P-Body Formation To Promote Viral RNA Synthesis. J Virol. 2021;95: e0092221. doi:10.1128/JVI.00922-21

33. Kanakamani S, Suresh PS, Venkatesh T. Regulation of processing bodies: From viruses to cancer epigenetic machinery. Cell Biol Int. John Wiley & Sons, Ltd; 2021;45: 708– 719. doi:10.1002/cbin.11527

34. Gaete-Argel A, Márquez CL, Barriga GP, Soto-Rifo R, Valiente-Echeverría F. Strategies for Success. Viral Infections and Membraneless Organelles. Front Cell Infect Microbiol. 2019;9: 336. doi:10.3389/fcimb.2019.00336

35. McCormick C, Khaperskyy DA. Translation inhibition and stress granules in the antiviral immune response. Nature Reviews Immunology. Nature Publishing Group; 2017;17: 647–660. doi:10.1038/nri.2017.63

36. Tsai W-C, Lloyd RE. Cytoplasmic RNA Granules and Viral Infection. Annu Rev Virol. Annual Reviews; 2014;1: 147–170. doi:10.1146/annurev-virology-031413-085505

37. Onomoto K, Jogi M, Yoo J-S, Narita R, Morimoto S, Takemura A, et al. Critical role of an antiviral stress granule containing RIG-I and PKR in viral detection and innate immunity. Kanai A, editor. PLoS ONE. Public Library of Science; 2012;7: e43031. doi:10.1371/journal.pone.0043031

38. Abernathy E, Gilbertson S, Alla R, Glaunsinger B. Viral Nucleases Induce an mRNA Degradation-Transcription Feedback Loop in Mammalian Cells. Cell Host and Microbe. 2015;18: 243–253. doi:10.1016/j.chom.2015.06.019

39. Núñez RD, Budt M, Saenger S, Paki K, Arnold U, Sadewasser A, et al. The RNA Helicase DDX6 Associates with RIG-I to Augment Induction of Antiviral Signaling. Int J Mol Sci. Multidisciplinary Digital Publishing Institute; 2018;19: 1877. doi:10.3390/ijms19071877

40. Balinsky CA, Schmeisser H, Wells AI, Ganesan S, Jin T, Singh K, et al. IRAV (FLJ11286), an Interferon-Stimulated Gene with Antiviral Activity against Dengue Virus, Interacts with MOV10. Diamond MS, editor. J Virol. 2017;91: 504. doi:10.1128/JVI.01606-16

41. Wang H, Chang L, Wang X, Su A, Feng C, Fu Y, et al. MOV10 interacts with Enterovirus 71 genomic 5’UTR and modulates viral replication. Biochem Biophys Res Commun. 2016;479: 571–577. doi:10.1016/j.bbrc.2016.09.112

42. Cuevas RA, Ghosh A, Wallerath C, Hornung V, Coyne CB, Sarkar SN. MOV10 Provides Antiviral Activity against RNA Viruses by Enhancing RIG-I-MAVS-Independent IFN Induction. J Immunol. 2016;196: 3877–3886. doi:10.4049/jimmunol.1501359

43. Burdick R, Smith JL, Chaipan C, Friew Y, Chen J, Venkatachari NJ, et al. P body-associated protein Mov10 inhibits HIV-1 replication at multiple stages. J Virol. 2010;84: 10241–10253. doi:10.1128/JVI.00585-10

44. Burgess HM, Mohr I. Cellular 5“-3” mRNA exonuclease Xrn1 controls double-stranded RNA accumulation and anti-viral responses. Cell Host and Microbe. 2015;17: 332–344. doi:10.1016/j.chom.2015.02.003

45. Lumb JH, Li Q, Popov LM, Ding S, Keith MT, Merrill BD, et al. DDX6 Represses Aberrant Activation of Interferon-Stimulated Genes. CellReports. ElsevierCompany; 2017;20: 819–831. doi:10.1016/j.celrep.2017.06.085

46. V’kovski P, Kratzel A, Steiner S, Stalder H, Thiel V. Coronavirus biology and replication: implications for SARS-CoV-2. Nat Rev Microbiol. Nature Publishing Group; 2020;5: 536–16. doi:10.1038/s41579-020-00468-6

47. Cui J, Li F, Shi Z-L. Origin and evolution of pathogenic coronaviruses. Springer US; 2019;: 1–12. doi:10.1038/s41579-018-0118-9

48. Banerjee A, Doxey AC, Tremblay BJ-M, Mansfield MJ, Subudhi S, Hirota JA, et al. Predicting the recombination potential of severe acute respiratory syndrome coronavirus 2 and Middle East respiratory syndrome coronavirus. J Gen Virol. Microbiology Society; 2020;101:<otherinfo> 12</otherinfo>51–1260. doi:10.1099/jgv.0.001491

49. Banerjee A, Doxey AC, Mossman K, Irving AT. Unraveling the Zoonotic Origin and Transmission of SARS-CoV-2. Trends Ecol Evol. 2021;36: 180–184. doi:10.1016/j.tree.2020.12.002

50. Zhou H, Ji J, Chen X, Bi Y, Li J, Wang Q, et al. Identification of novel bat coronaviruses sheds light on the evolutionary origins of SARS-CoV-2 and related viruses. Cell. 2021;184: 4380–4391.e14. doi:10.1016/j.cell.2021.06.008

51. Li W, Shi Z, Yu M, Ren W, Smith C, Epstein JH, et al. Bats are natural reservoirs of SARS-like coronaviruses. Science. American Association for the Advancement of Science; 2005;310: 676–679. doi:10.1126/science.1118391

52. Fielding CA, Jones GW, McLoughlin RM, McLeod L, Hammond VJ, Uceda J, et al. Interleukin-6 signaling drives fibrosis in unresolved inflammation. Immunity. 2014;40: 40–50. doi:10.1016/j.immuni.2013.10.022

53. Blanco-Melo D, tenover BR. Imbalanced host response to SARS-CoV-2 drives development of COVID-19. Cell. 2020;: 1–46. doi:10.1016/j.cell.2020.04.026

54. Pedersen SF, Ho Y-C. SARS-CoV-2: a storm is raging. Journal of Clinical Investigation. 2020;130: 2202–2205. doi:10.1172/JCI137647

55. Zou L, Ruan F, Huang M, Liang L, Huang H, Hong Z, et al. SARS-CoV-2 Viral Load in Upper Respiratory Specimens of Infected Patients. N Engl J Med. Massachusetts Medical Society; 2020;382: 1177–1179. doi:10.1056/NEJMc2001737

56. Libby P, Lüscher T. COVID-19 is, in the end, an endothelial disease. Eur Heart J. 3rd ed. 2020;41: 3038–3044. doi:10.1093/eurheartj/ehaa623

57. Chen G, Wu D, Guo W, Cao Y, Huang D, Wang H, et al. Clinical and immunological features of severe and moderate coronavirus disease 2019. J Clin Invest. American Society for Clinical Investigation; 2020;130: 2620–2629. doi:10.1172/JCI137244

58. Pons S, Fodil S, Azoulay E, Zafrani L. The vascular endothelium: the cornerstone of organ dysfunction in severe SARS-CoV-2 infection. Crit Care. BioMed Central; 2020;24: 353–8. doi:10.1186/s13054-020-03062-7

59. Lowenstein CJ, Solomon SD. Severe COVID-19 is a Microvascular Disease. Circulation. Lippincott Williams & Wilkins Hagerstown, MD; 2020;191: 148. doi:10.1161/CIRCULATIONAHA.120.050354

60. Varga Z, Flammer AJ, Steiger P, Haberecker M, Andermatt R, Zinkernagel AS, et al. Endothelial cell infection and endotheliitis in COVID-19. The Lancet. Elsevier Ltd; 2020;395: 1417–1418. doi:10.1016/S0140-6736(20)30937-5

61. Amraei R, Rahimi N. COVID-19, Renin-Angiotensin System and Endothelial Dysfunction. Cells. Multidisciplinary Digital Publishing Institute; 2020;9: 1652. doi:10.3390/cells9071652

62. Teuwen L-A, Geldhof V, Pasut A, Carmeliet P. COVID-19: the vasculature unleashed. Nature Reviews Immunology. Nature Publishing Group; 2020;20: 389–391. doi:10.1038/s41577-020-0343-0

63. Gu SX, Tyagi T, Jain K, Gu VW, Lee SH, Hwa JM, et al. Thrombocytopathy and endotheliopathy: crucial contributors to COVID-19 thromboinflammation. Nat Rev Cardiol. Nature Publishing Group; 2021;18: 194–209. doi:10.1038/s41569-020-00469-1

64. Suvorava T, Kaesemeyer W. Targeting the Vascular Endothelium in the Treatment of COVID-19. J Cardiovasc Pharmacol. 2021;77: 1–3. doi:10.1097/FJC.0000000000000932

65. Bastard P, Rosen LB, Zhang Q, Michailidis E, Hoffmann H-H, Zhang Y, et al. Auto-antibodies against type I IFNs in patients with life-threatening COVID-19. Science. 2020;129: eabd4585. doi:10.1126/science.abd4585

66. Zhang Q, Bastard P, Liu Z, Le Pen J, Moncada-Velez M, Chen J, et al. Inborn errors of type I IFN immunity in patients with life-threatening COVID-19. Science. 2020;: eabd4570. doi:10.1126/science.abd4570

67. Park A, Iwasaki A. Type I and Type III Interferons - Induction, Signaling, Evasion, and Application to Combat COVID-19. Cell Host and Microbe. 2020;27: 870–878. doi:10.1016/j.chom.2020.05.008

68. Thorne LG, Bouhaddou M, Reuschl A-K, Zuliani-Alvarez L, Polacco B, Pelin A, et al. Evolution of enhanced innate immune evasion by the SARS-CoV-2 B.1.1.7 UK variant. PlosPath. Cold Spring Harbor Laboratory; 2021;: 2021.06.06.446826. doi:10.1101/2021.06.06.446826

69. Sposito B, Broggi A, Pandolfi L, Crotta S, Clementi N, Ferrarese R, et al. The interferon landscape along the respiratory tract impacts the severity of COVID-19. Cell. 2021;184: 4953–4968.e16. doi:10.1016/j.cell.2021.08.016

70. Ziegler CGK, Miao VN, Owings AH, Navia AW, Tang Y, Bromley JD, et al. Impaired local intrinsic immunity to SARS-CoV-2 infection in severe COVID-19. Cell. 2021;184: 4713–4733.e22. doi:10.1016/j.cell.2021.07.023

71. Zhou Z, Ren L, Zhang L, Zhong J, Xiao Y, Jia Z, et al. Heightened Innate Immune Responses in the Respiratory Tract of COVID-19 Patients. Cell Host and Microbe. 2020;27: 883–890.e2. doi:10.1016/j.chom.2020.04.017

72. Hadjadj J, Yatim N, Barnabei L, Corneau A, Boussier J, Smith N, et al. Impaired type I interferon activity and inflammatory responses in severe COVID-19 patients. Science. 2020;369: 718–724. doi:10.1126/science.abc6027

73. Banerjee A, El-Sayes N, Budylowski P, Jacob RA, Richard D, Maan H, et al. Experimental and natural evidence of SARS-CoV-2-infection-induced activation of type I interferon responses. iScience. 2021;24: 102477. doi:10.1016/j.isci.2021.102477

74. Yuen C-K, Lam J-Y, Wong W-M, Mak L-F, Wang X, Chu H, et al. SARS-CoV-2 nsp13, nsp14, nsp15 and orf6 function as potent interferon antagonists. Emerging Microbes & Infections. Taylor & Francis; 2020;9: 1418–1428. doi:10.1080/22221751.2020.1780953

75. Miorin L, Kehrer T, Sanchez-Aparicio MT, Zhang K, Cohen P, Patel RS, et al. SARS-CoV-2 Orf6 hijacks Nup98 to block STAT nuclear import and antagonize interferon signaling. Proc Natl Acad Sci USA. 2020;5: 202016650–11. doi:10.1073/pnas.2016650117

76. Banerjee AK, Blanco MR, Bruce EA, Honson DD, Chen LM, Chow A, et al. SARS-CoV-2 disrupts splicing, translation, and protein trafficking to suppress host defenses. Cell. Elsevier Inc; 2020;: 1–66. doi:10.1016/j.cell.2020.10.004

77. Konno Y, Kimura I, Uriu K, Fukushi M, Irie T, Koyanagi Y, et al. SARS-CoV-2 ORF3b Is a Potent Interferon Antagonist Whose Activity Is Increased by a Naturally Occurring Elongation Variant. CellReports. 2020;32: 108185. doi:10.1016/j.celrep.2020.108185

78. Yin X, Riva L, Pu Y, Martin-Sancho L, Kanamune J, Yamamoto Y, et al. MDA5 Governs the Innate Immune Response to SARS-CoV-2 in Lung Epithelial Cells. CellReports. 2021;34: 108628. doi:10.1016/j.celrep.2020.108628

79. Lei X, Dong X, Ma R, Wang W, Xiao X, Tian Z, et al. Activation and evasion of type I interferon responses by SARS-CoV-2. Nature Communications. Nature Publishing Group; 2020;11: 3810–12. doi:10.1038/s41467-020-17665-9

80. Beyer DK, Forero A. Mechanisms of Antiviral Immune Evasion of SARS-CoV-2. J Mol Biol. 2021;: 167265. doi:10.1016/j.jmb.2021.167265

81. Xia H, Cao Z, Xie X, Zhang X, Chen JY-C, Wang H, et al. Evasion of Type I Interferon by SARS-CoV-2. CellReports. 2020;33: 108234. doi:10.1016/j.celrep.2020.108234

82. Suryawanshi RK, Koganti R, Agelidis A, Patil CD, Shukla D. Dysregulation of Cell Signaling by SARS-CoV-2. Trends in Microbiology. 2021;29: 224–237. doi:10.1016/j.tim.2020.12.007

83. Wickenhagen A, Sugrue E, Lytras S, Kuchi S, Noerenberg M, Turnbull ML, et al. A prenylated dsRNA sensor protects against severe COVID-19. Science. American Association for the Advancement of Science; 2021;: eabj3624. doi:10.1126/science.abj3624

84. Hsu JC-C, Laurent-Rolle M, Pawlak JB, Wilen CB, Cresswell P. Translational shutdown and evasion of the innate immune response by SARS-CoV-2 NSP14 protein. Proc Natl Acad Sci USA. National Academy of Sciences; 2021;118. doi:10.1073/pnas.2101161118

85. Zhao Y, Sui L, Wu P, Wang W, Wang Z, Yu Y, et al. A dual-role of SARS-CoV-2 nucleocapsid protein in regulating innate immune response. Signal Transduct Target Ther. Nature Publishing Group; 2021;6: 331–14. doi:10.1038/s41392-021-00742-w

86. Sola I, Galán C, Mateos-Gómez PA, Palacio L, Zuñiga S, Cruz JL, et al. The polypyrimidine tract-binding protein affects coronavirus RNA accumulation levels and relocalizes viral RNAs to novel cytoplasmic domains different from replication-transcription sites. J Virol. 3rd ed. 2011;85: 5136–5149. doi:10.1128/JVI.00195-11

87. Raaben M, Groot Koerkamp MJA, Rottier PJM, de Haan CAM. Mouse hepatitis coronavirus replication induces host translational shutoff and mRNA decay, with concomitant formation of stress granules and processing bodies. Cellular Microbiology. 2007;9: 2218–2229. doi:10.1111/j.1462-5822.2007.00951.x

88. Ackermann M, Mentzer SJ, Kolb M, Jonigk D. Inflammation and Intussusceptive Angiogenesis in COVID-19: everything in and out of Flow. Eur Respir J. 2020;383: 2003147. doi:10.1183/13993003.03147-2020

89. Nascimento Conde J, Schutt WR, Gorbunova EE, Mackow ER. Recombinant ACE2 Expression Is Required for SARS-CoV-2 To Infect Primary Human Endothelial Cells and Induce Inflammatory and Procoagulative Responses. Patton JT, editor. mBio. American Society for Microbiology; 2020;11. doi:10.1128/mBio.03185-20

90. Banerjee A, Nasir JA, Budylowski P, Yip L, Aftanas P, Christie N, et al. Isolation, Sequence, Infectivity, and Replication Kinetics of Severe Acute Respiratory Syndrome Coronavirus 2. Emerging Infect Dis. 2020;26: 2054–2063. doi:10.3201/eid2609.201495

91. Castle EL, Robinson C-A, Douglas P, Rinker KD, Corcoran JA. Viral manipulation of a mechanoresponsive signaling axis disassembles processing bodies. Mol Cell Biol. 2021;: MCB0039921. doi:10.1128/MCB.00399-21

92. Singh J, Pandit P, McArthur AG, Banerjee A, Mossman K. Evolutionary trajectory of SARS-CoV-2 and emerging variants. Virology Journal. BioMed Central; 2021;18: 166– 21. doi:10.1186/s12985-021-01633-w

93. Harvey WT, Carabelli AM, Jackson B, Gupta RK, Thomson EC, Harrison EM, et al. SARS-CoV-2 variants, spike mutations and immune escape. Nat Rev Microbiol. Nature Publishing Group; 2021;19: 409–424. doi:10.1038/s41579-021-00573-0

94. Sola I, Almazán F, Zuñiga S, Enjuanes L. Continuous and Discontinuous RNA Synthesis in Coronaviruses. Annu Rev Virol. Annual Reviews; 2015;2: 265–288. doi:10.1146/annurev-virology-100114-055218

95. Ayache J, Bénard M, Ernoult-Lange M, Minshall N, Standart N, Kress M, et al. P-body assembly requires DDX6 repression complexes rather than decay or Ataxin2/2L complexes. Matera AG, editor. Mol Biol Cell. The American Society for Cell Biology; 2015;26: 2579–2595. doi:10.1091/mbc.E15-03-0136

96. Gordon DE, Jang GM, Bouhaddou M, Xu J, Obernier K, White KM, et al. A SARS-CoV-2 protein interaction map reveals targets for drug repurposing. Nature. Nature Publishing Group; 2020;583: 459–468. doi:10.1038/s41586-020-2286-9

97. Kleer M, MacNeil G, Adam N, Pringle ES, Corcoran JA. A panel of KSHV mutants in the polycistronic kaposin locus for precise analysis of individual protein products. J Virol. American Society for Microbiology 1752 N St., N.W., Washington, DC; 2021;: JVI0156021. doi:10.1128/JVI.01560-21

98. Robinson C-A, Singh GK, Kleer M, Castle EL, Boudreau BQ, Corcoran JA. Kaposi’s sarcoma-associated herpesvirus (KSHV) utilizes the NDP52/CALCOCO2 selective autophagy receptor to disassemble processing bodies. PLoS Pathogens. 2022;: 1–61. doi:10.1101/2021.02.07.430164

99. Min Y-Q, Mo Q, Wang J, Deng F, Wang H, Ning Y-J. SARS-CoV-2 nsp1: Bioinformatics, Potential Structural and Functional Features, and Implications for Drug/Vaccine Designs. Front Microbiol. Frontiers; 2020;11: 587317. doi:10.3389/fmicb.2020.587317

100. Zeng W, Liu G, Ma H, Zhao D, Yang Y, Liu M, et al. Biochemical characterization of SARS-CoV-2 nucleocapsid protein. Biochem Biophys Res Commun. 2020;527: 618– 623. doi:10.1016/j.bbrc.2020.04.136

101. Kedersha N, Stoecklin G, Ayodele M, Yacono P, Lykke-Andersen J, Fritzler MJ, et al. Stress granules and processing bodies are dynamically linked sites of mRNP remodeling. J Cell Biol. 2005;169: 871–884. doi:10.1083/jcb.200502088

102. Bai Z, Cao Y, Liu W, Li J. The SARS-CoV-2 Nucleocapsid Protein and Its Role in Viral Structure, Biological Functions, and a Potential Target for Drug or Vaccine Mitigation. Viruses. Multidisciplinary Digital Publishing Institute; 2021;13: 1115. doi:10.3390/v13061115

103. Peng T-Y, Lee K-R, Tarn W-Y. Phosphorylation of the arginine/serine dipeptide-rich motif of the severe acute respiratory syndrome coronavirus nucleocapsid protein modulates its multimerization, translation inhibitory activity and cellular localization. FEBS J. John Wiley & Sons, Ltd; 2008;275: 4152–4163. doi:10.1111/j.1742-4658.2008.06564.x

104. McBride R, van Zyl M, Fielding BC. The coronavirus nucleocapsid is a multifunctional protein. Viruses. Multidisciplinary Digital Publishing Institute; 2014;6: 2991–3018. doi:10.3390/v6082991

105. Cong Y, Ulasli M, Schepers H, Mauthe M, V’kovski P, Kriegenburg F, et al. Nucleocapsid Protein Recruitment to Replication-Transcription Complexes Plays a Crucial Role in Coronaviral Life Cycle. Dutch RE, editor. J Virol. American Society for Microbiology Journals; 2020;94: 181. doi:10.1128/JVI.01925-19

106. Lu S, Ye Q, Singh D, Villa E, Cleveland DW, Corbett KD. The SARS-CoV-2 Nucleocapsid phosphoprotein forms mutually exclusive condensates with RNA and the membrane-associated M protein. PlosPath. 2020;18: 479. doi:10.1101/2020.07.30.228023

107. Cascarina SM, Ross ED. A proposed role for the SARS-CoV-2 nucleocapsid protein in the formation and regulation of biomolecular condensates. FASEB j. 2020;34: 9832– 9842. doi:10.1096/fj.202001351

108. Chen H, Cui Y, Han X, Hu W, Sun M, Zhang Y, et al. Liquid-liquid phase separation by SARS-CoV-2 nucleocapsid protein and RNA. Cell Res. Nature Publishing Group; 2020;30: 1143–1145. doi:10.1038/s41422-020-00408-2

109. Savastano A, Ibáñez de Opakua A, Rankovic M, Zweckstetter M. Nucleocapsid protein of SARS-CoV-2 phase separates into RNA-rich polymerase-containing condensates. Nature Communications. Nature Publishing Group; 2020;11: 6041–10. doi:10.1038/s41467-020-19843-1

110. Iserman C, Roden CA, Boerneke MA, Sealfon RSG, McLaughlin GA, Jungreis I, et al. Genomic RNA Elements Drive Phase Separation of the SARS-CoV-2 Nucleocapsid. Mol Cell. 2020;80: 1078–1091.e6. doi:10.1016/j.molcel.2020.11.041

111. Carlson CR, Asfaha JB, Ghent CM, Howard CJ, Hartooni N, Safari M, et al. Phosphoregulation of Phase Separation by the SARS-CoV-2 N Protein Suggests a Biophysical Basis for its Dual Functions. Mol Cell. 2020;80: 1092–1103.e4. doi:10.1016/j.molcel.2020.11.025

112. Cubuk J, Alston JJ, Incicco JJ, Singh S, Stuchell-Brereton MD, Ward MD, et al. The SARS-CoV-2 nucleocapsid protein is dynamic, disordered, and phase separates with RNA. Nature Communications. Nature Publishing Group; 2021;12: 1936–17. doi:10.1038/s41467-021-21953-3

113. Sa Ribero M, Jouvenet N, Dreux M, Nisole S. Interplay between SARS-CoV-2 and the type I interferon response. PLoS Pathogens. Public Library of Science; 2020;16: e1008737. doi:10.1371/journal.ppat.1008737

114. Gori Savellini G, Anichini G, Gandolfo C, Cusi MG. SARS-CoV-2 N Protein Targets TRIM25-Mediated RIG-I Activation to Suppress Innate Immunity. Viruses. Multidisciplinary Digital Publishing Institute; 2021;13: 1439. doi:10.3390/v13081439

115. Oliveira SC, de Magalhães MTQ, Homan EJ. Immunoinformatic Analysis of SARS-CoV-2 Nucleocapsid Protein and Identification of COVID-19 Vaccine Targets. Front Immunol. Frontiers; 2020;11: 587615. doi:10.3389/fimmu.2020.587615

116. Chan JF-W, Kok K-H, Zhu Z, Chu H, To KK-W, Yuan S, et al. Genomic characterization of the 2019 novel human-pathogenic coronavirus isolated from a patient with atypical pneumonia after visiting Wuhan. Emerging Microbes & Infections. 2020;9: 221–236. doi:10.1080/22221751.2020.1719902

117. Parker MD, Lindsey BB, Leary S, Gaudieri S, Chopra A, Wyles M, et al. Subgenomic RNA identification in SARS-CoV-2 genomic sequencing data. Genome Res. Cold Spring Harbor Lab; 2021;31: 645–658. doi:10.1101/gr.268110.120

118. Leary S, Gaudieri S, Parker MD, Chopra A, James I, Pakala S, et al. Generation of a Novel SARS-CoV-2 Sub-genomic RNA Due to the R203K/G204R Variant in Nucleocapsid: Homologous Recombination has Potential to Change SARS-CoV-2 at Both Protein and RNA Level. Pathog Immun. 2021;6: 27–49. doi:10.20411/pai.v6i2.460

119. Saffran HA, Pare JM, Corcoran JA, Weller SK, Smiley JR. Herpes simplex virus eliminates host mitochondrial DNA. EMBO Rep. 2007;8: 188–193. doi:10.1038/sj.embor.7400878

120. Corcoran JA, Saffran HA, Duguay BA, Smiley JR. Herpes simplex virus UL12.5 targets mitochondria through a mitochondrial localization sequence proximal to the N terminus. J Virol. 2009;83: 2601–2610. doi:10.1128/JVI.02087-08

121. Gordon DE, Hiatt J, Bouhaddou M, Rezelj VV, Ulferts S, Braberg H, et al. Comparative host-coronavirus protein interaction networks reveal pan-viral disease mechanisms. Science. 2020;11: eabe9403–38. doi:10.1126/science.abe9403

122. Nabeel-Shah S, Lee H, Ahmed N, Burke GL, Farhangmehr S, Ashraf K, et al. SARS-CoV-2 nucleocapsid protein binds host mRNAs and attenuates stress granules to impair host stress response. iScience. 2022;25: 103562. doi:10.1016/j.isci.2021.103562

123. Luo L, Li Z, Zhao T, Ju X, Ma P, Jin B, et al. SARS-CoV-2 nucleocapsid protein phase separates with G3BPs to disassemble stress granules and facilitate viral production. Sci Bull (Beijing). 2021;66: 1194–1204. doi:10.1016/j.scib.2021.01.013

124. Zheng Z-Q, Wang S-Y, Xu Z-S, Fu Y-Z, Wang Y-Y. SARS-CoV-2 nucleocapsid protein impairs stress granule formation to promote viral replication. Cell Discov. Nature Publishing Group; 2021;7: 38–11. doi:10.1038/s41421-021-00275-0

125. Wang J, Shi C, Xu Q, Yin H. SARS-CoV-2 nucleocapsid protein undergoes liquid-liquid phase separation into stress granules through its N-terminal intrinsically disordered region. Cell Discov. Nature Publishing Group; 2021;7: 5–5. doi:10.1038/s41421-020-00240-3

126. Dougherty JD, Reineke LC, Lloyd RE. mRNA decapping enzyme 1a (Dcp1a)-induced translational arrest through protein kinase R (PKR) activation requires the N-terminal enabled vasodilator-stimulated protein homology 1 (EVH1) domain. J Biol Chem. 2014;289: 3936–3949. doi:10.1074/jbc.M113.518191

127. Schmidt N, Lareau CA, Keshishian H, Ganskih S, Schneider C, Hennig T, et al. The SARS-CoV-2 RNA-protein interactome in infected human cells. Nat Microbiol. Nature Publishing Group; 2021;6: 339–353. doi:10.1038/s41564-020-00846-z

128. Tibble RW, Depaix A, Kowalska J, Jemielity J, Gross JD. Biomolecular condensates amplify mRNA decapping by coupling protein interactions with conformational changes in Dcp1/Dcp2. 2020;60th Anniversary: 600–36. doi:10.1101/2020.07.09.195057

129. Tenekeci U, Poppe M, Beuerlein K, Buro C, Müller H, Weiser H, et al. K63-Ubiquitylation and TRAF6 Pathways Regulate Mammalian P-Body Formation and mRNA Decapping. Mol Cell. 2016;62: 943–957. doi:10.1016/j.molcel.2016.05.017

130. Li Y, Chen R, Zhou Q, Xu Z, Li C, Wang S, et al. LSm14A is a processing body-associated sensor of viral nucleic acids that initiates cellular antiviral response in the early phase of viral infection. Proc Natl Acad Sci USA. 2012;109: 11770–11775. doi:10.1073/pnas.1203405109

131. Rzeczkowski K, Beuerlein K, Müller H, Dittrich-Breiholz O, Schneider H, Kettner-Buhrow D, et al. c-Jun N-terminal kinase phosphorylates DCP1a to control formation of P bodies. J Cell Biol. 2011;194: 581–596. doi:10.1083/jcb.201006089

132. Stoecklin G, Mayo T, Anderson P. ARE-mRNA degradation requires the 5“-3” decay pathway. EMBO Rep. 2006;7: 72–77. doi:10.1038/sj.embor.7400572

133. Mendoza EJ, Manguiat K, Wood H, Drebot M. Two Detailed Plaque Assay Protocols for the Quantification of Infectious SARS-CoV-2. Curr Protoc Microbiol. John Wiley & Sons, Ltd; 2020;57: cpmc105. doi:10.1002/cpmc.105

134. Kamentsky L, Jones TR, Fraser A, Bray M-A, Logan DJ, Madden KL, et al. Improved structure, function and compatibility for CellProfiler: modular high-throughput image analysis software. Bioinformatics. 2011;27: 1179–1180. doi:10.1093/bioinformatics/btr095

135. Johnston BP, Pringle ES, McCormick C. KSHV activates unfolded protein response sensors but suppresses downstream transcriptional responses to support lytic replication. Swaminathan S, editor. PLoS Pathogens. 2019;15: e1008185. doi:10.1371/journal.ppat.1008185

