## Supplementary figures and images for "Human coronaviruses disassemble processing bodies"

### Supplemental Figures

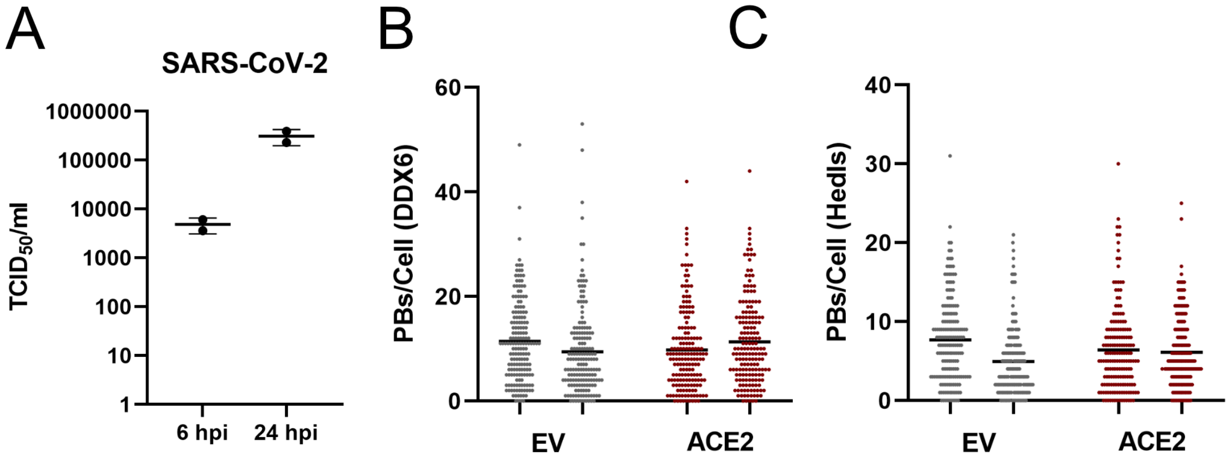

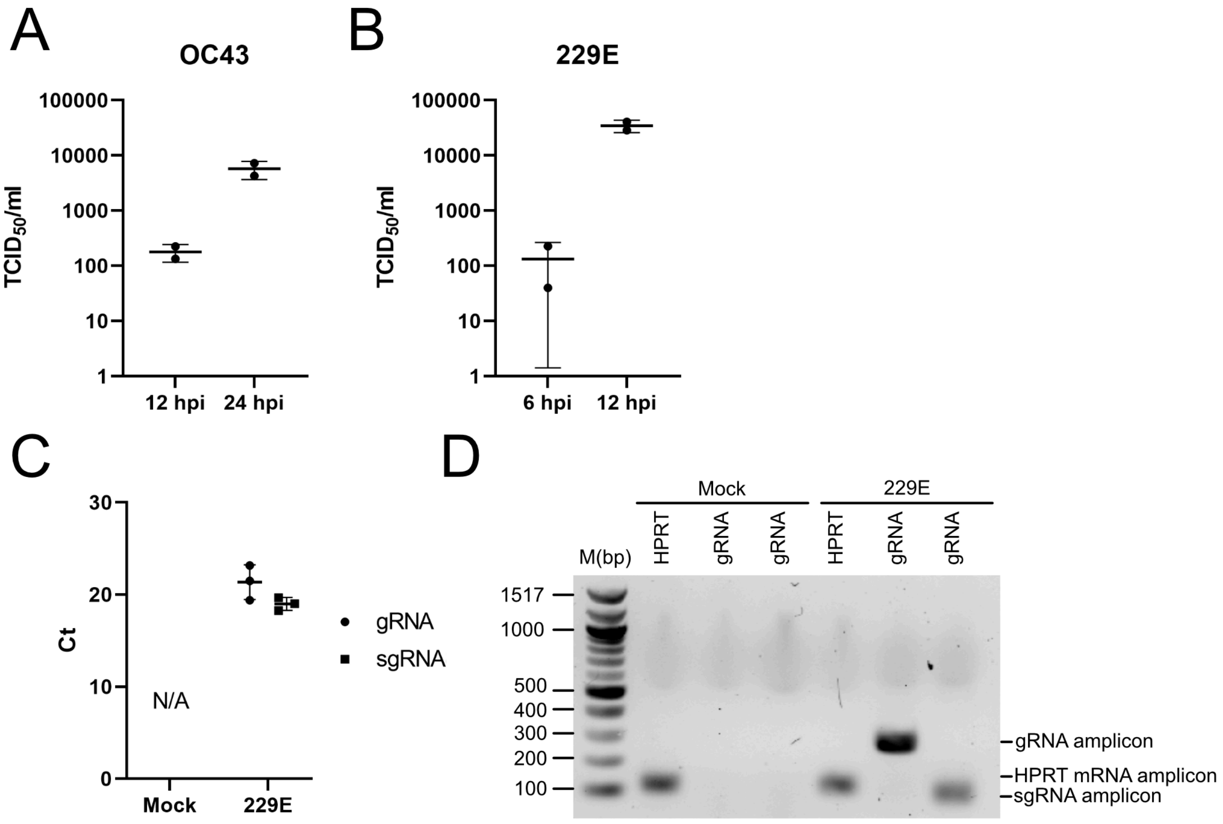

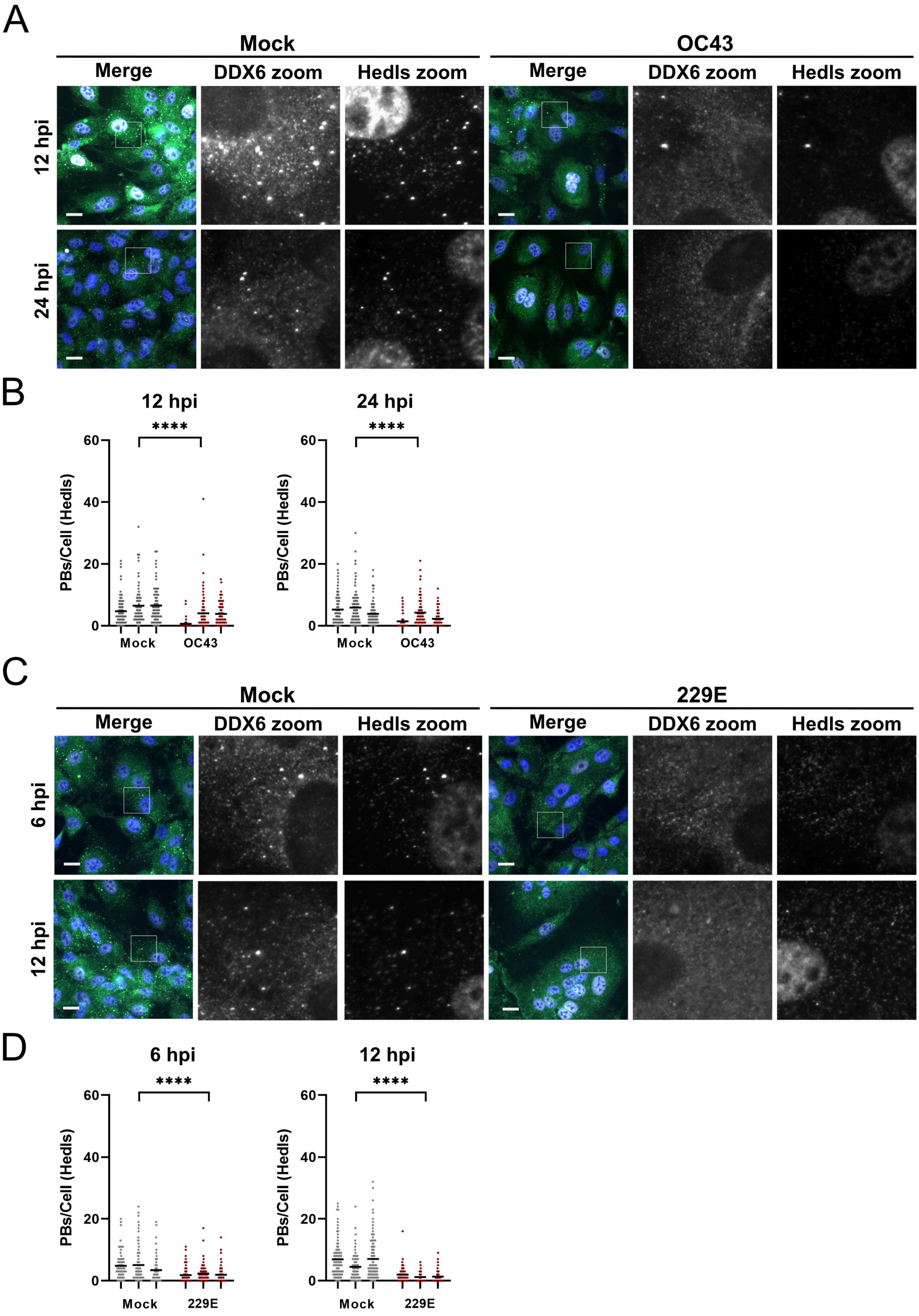

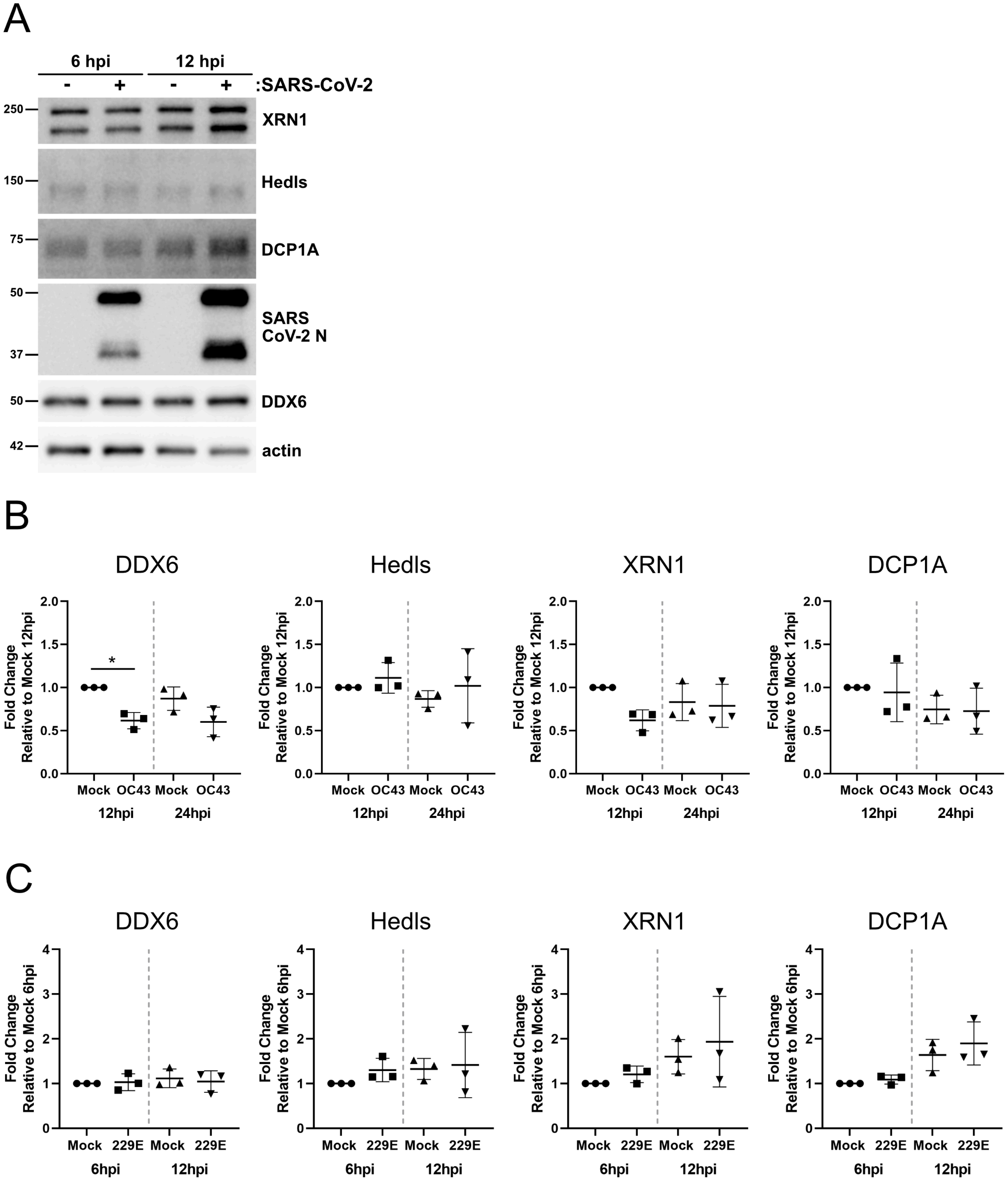

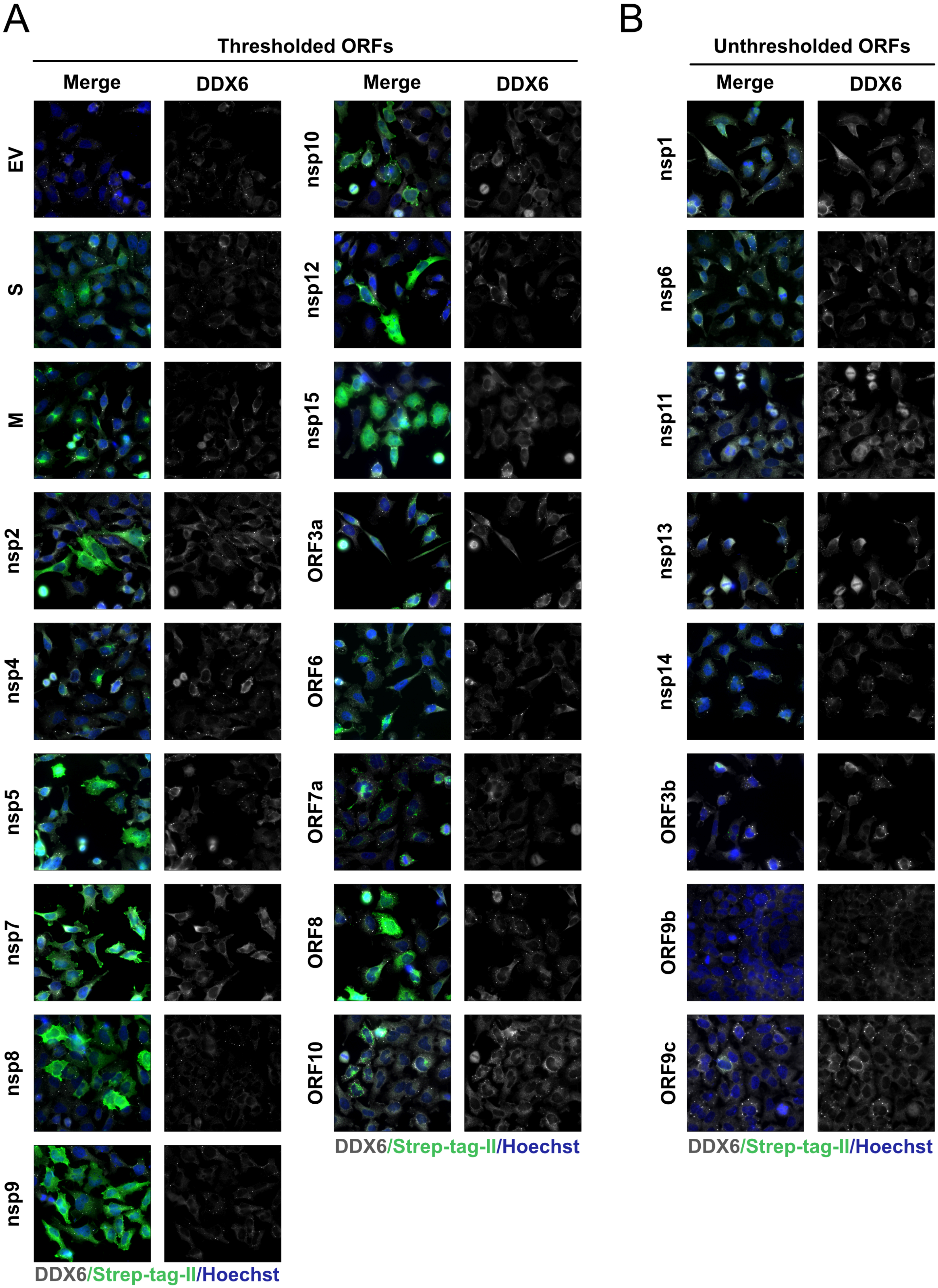

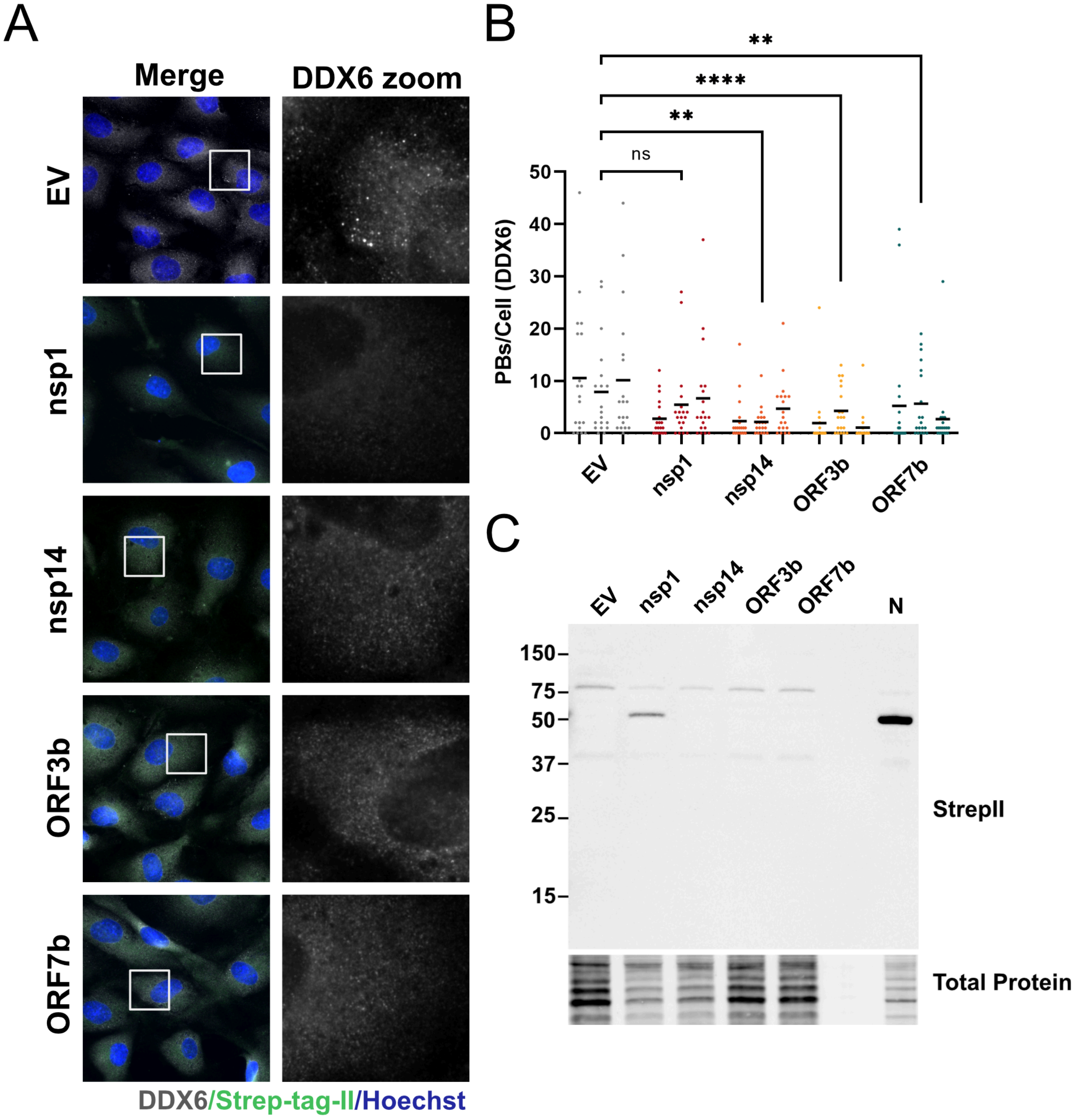

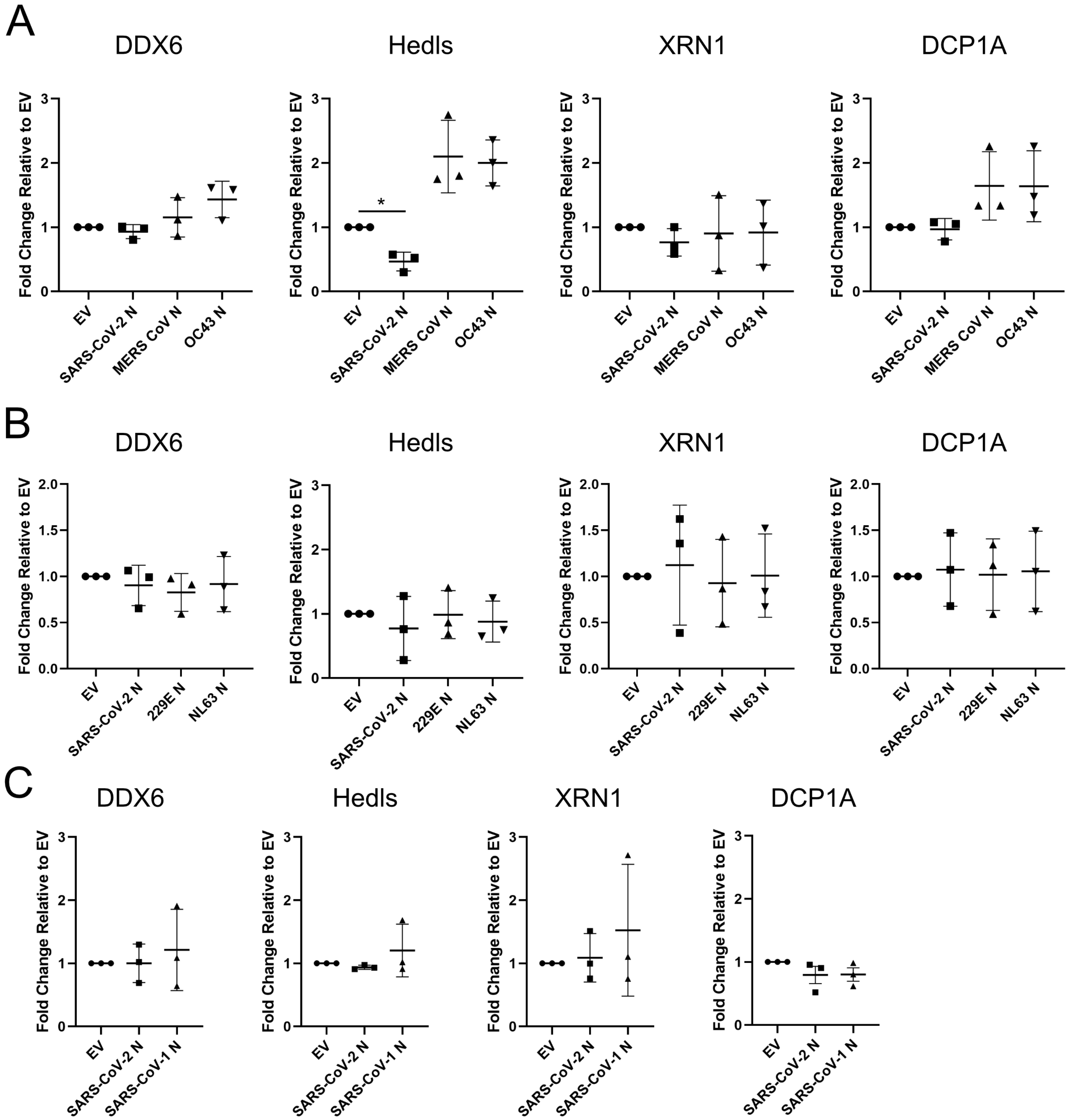

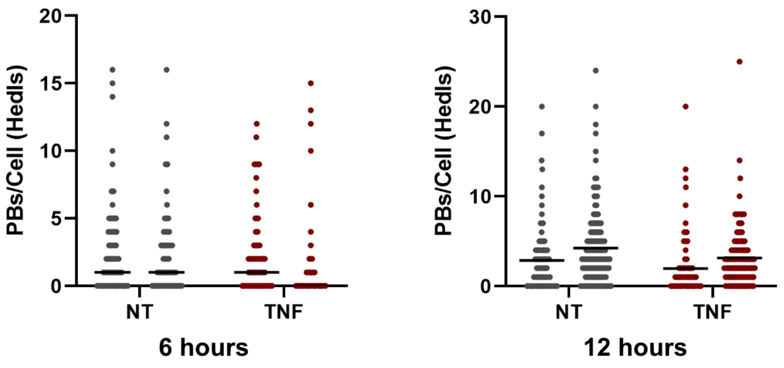
